# Helicase family proteins promote antiviral ADP-ribosylation of DNA purines

**DOI:** 10.64898/2026.11.02.763693

**Authors:** Trevor Griesman, Adini Q. Arifah, Eric J. Tomko, Yongjun Tan, Ning Tsao, Jackson T. Trotter, Sharvika S. Khochare, Nico A. Carlson, Nima Mosammaparast, Dapeng Zhang, Eric A. Galburt, Constantinos Patinios, Michele LeRoux

**Affiliations:** Department of Molecular Microbiology, Washington University in Saint Louis, St. Louis, MO, USA; LSC-EMBL Partnership Institute for Genome Editing Technologies, Life Sciences Center, Vilnius University, Vilnius, Lithuania; Department of Biochemistry and Molecular Biophysics, Washington University in Saint Louis, St. Louis, MO, USA; Department of Biology, Saint Louis University, St. Louis, MO, USA; Department of Pathology and Immunology, Washington University in Saint Louis, St. Louis, MO, USA

## Abstract

DarTG defense systems studied to date contain a DarT1 or DarT2 toxin that blocks phage replication by ADP-ribosylating DNA and a NADAR or macrodomain antitoxin, DarG, that reverses this modification at homeostasis. Here, we identify additional DarTG subfamilies, one of which comprises a DarT3 toxin, an ADP-ribosylhydrolase (ARH)-domain antitoxin, and a helicase family protein, DarH. The *Escherichia coli* DarHTG3 system confers broad phage defense, requires DarH and, unlike previously characterized DarTG systems, modifies adenine. Characterization of additional DarH-containing DarTG systems reveals that a subset also target guanine. Our data support a model in which ADP-bound DarH functions as a processivity factor by recruiting DarT to its single-stranded DNA substrate and allowing for ADP-ribosylation of multiple target sites. These findings reveal how a helicase family protein regulates an antiviral effector by promoting DNA engagement rather than unwinding and thereby expanding the functional repertoire of helicase family proteins in bacterial immunity.

## Introduction

Bacteria and their phages are locked in a rapidly evolving molecular arms race. The strong selective pressure from lytic viral infections drives ongoing innovation in bacterial defenses, and corresponding co-evolution of viral counter-defenses. One such defense, DarTG systems, are typically two gene toxin-antitoxin operons, in which the DarT toxin ADP-ribosylates single-stranded DNA (ssDNA), while the DarG antitoxin, an ADP-ribosylglycohydrolase, enzymatically removes this modification in uninfected cells^1–3^. Upon phage infection and through a yet unknown trigger, the activity of the toxin is unleashed, resulting in an accumulation of toxic DNA ADP-ribosylation that blocks DNA synthesis and transcription to prevent phage replication^4^. Two DarTG subfamilies have been reported, distinguished by their antitoxin and base target. DarTG1 systems carry a toxin that modifies guanine, which is reversed by an NAD-and Ribose-Associated (NADAR) class antitoxin, DarG1 (which we refer to as DarG_N_, for reasons detailed below)^2^. In contrast, DarT2 ADP-ribosylates thymine, and DarG2 (from this point, DarG_M_) is a macrodomain family-containing antitoxin^1,3^. The two systems are distinct, as the antitoxins are specific for their cognate toxin, and they defend against different phages^2,4^.

Here, we report the discovery of additional DarTG subfamilies, which carry a third, unrelated DarG from the ADP-ribosylhydrolase (ARH) family (which we term DarG_R_)^5^. These new DarTG subfamilies also encode a third gene in the operon, a predicted helicase family protein (which we term DarH). Helicase family proteins have been incorporated into antiviral defenses across domains of life^6–8^. Some, exemplified by the eukaryotic Retinoic acid Inducible Gene-I (RIG-I), lack classical helicase activity such as DNA unwinding or translocation, and act instead as sensors of viral nucleic acid^9^. More recent analyses have uncovered helicase family proteins incorporated into bacterial defense systems such as Hachiman, Gabija, Hna, and Druantia^6–8,10–12^. While some have been shown to be sensors of infection or effector proteins, the activities of most others, including DarH, remain unclear.

We performed an in-depth characterization of a DarHTG system from *E. coli* NILS37:A, which we refer to as EcoDarHT3G_R_. We find that this system displays robust phage defense against a broad set of phages. Further, unlike previously studied DarTG systems, EcoDarHT3G_R_ modifies the adenine nucleobase in ssDNA. The helicase family component, EcoDarH, strongly promotes DarT3 toxicity *in vivo* and is required for phage defense. We also found that DarH increases DarT activity *in vitro*, and unexpectedly, this does not occur via canonical helicase activity such as ATP-powered translocation and DNA unwinding. Instead, the ADP-bound form of EcoDarH enables EcoDarT3 to bind to ssDNA. These data reveal the mechanism by which an immune-associated helicase family protein regulates the activity of an antiviral effector, and extend the breadth of known prokaryotic DNA ADP-ribosylation.

## Results

### DarHTG3 is a phage defense system associated with a helicase family protein

DarTG systems were originally described as two-gene toxin-antitoxin systems^1,4^. To identify additional variants of these operons, we performed an initial gene-neighborhood analysis based on *E. coli* DarT1 and DarT2 (Extended Data Fig. 1)^13^. We discovered that a subset of *darT1*-like operons included an upstream gene, a predicted *uvrD*-class, superfamily-1 helicase, *darH* (Fig. 1a,b, Extended Data Fig. 2a). To systematically explore the prevalence of such systems and their relationship to known DarTGs, we performed a clustering analysis. Using DarT1 and DarT2 examples, we ran iterative Position Specific Iterated (PSI)-BLAST searches against the NCBI non-redundant database to identify DarT homologs and used sequence similarity-based network analysis to cluster them into groups^14,15^. Gene neighborhood analysis revealed two groups that were associated with NADAR or macrodomain proteins, corresponding to the characterized DarT1G_N_ and DarT2G_M_ subfamilies, respectively. However, we also identified three distinct DarT clusters, all of which were associated with a *darH* gene (1377 instances). In these systems, the downstream *darG* was either an ARH (1248 instances) or NADAR (129 instances). Group 3 clustered separately from either DarT1 or DarT2, and was found in association with only ARH DarGs. Groups 4 and 5 clustered closer to DarT1, and were associated with either DarG_R_ or DarG_N_. Previously, it was thought that each DarT type was always associated with the same antitoxin type (i.e. DarT1 with DarG_N_, DarT2 with DarG_M_), thus DarTs were numbered along with DarGs. However, our analysis reveals that DarT proteins from different subfamilies can share a DarG type. We therefore propose a naming convention where DarTs are numbered by group (DarT1-T5), and DarGs are designated by protein class (<u>N</u>ADAR = DarG_N_, <u>m</u>acrodomain = DarG_M_, A<u>R</u>H = DarG_R_). DarHT3G_R_ was independently identified while this work was in progress^16^.

**Figure 1:**
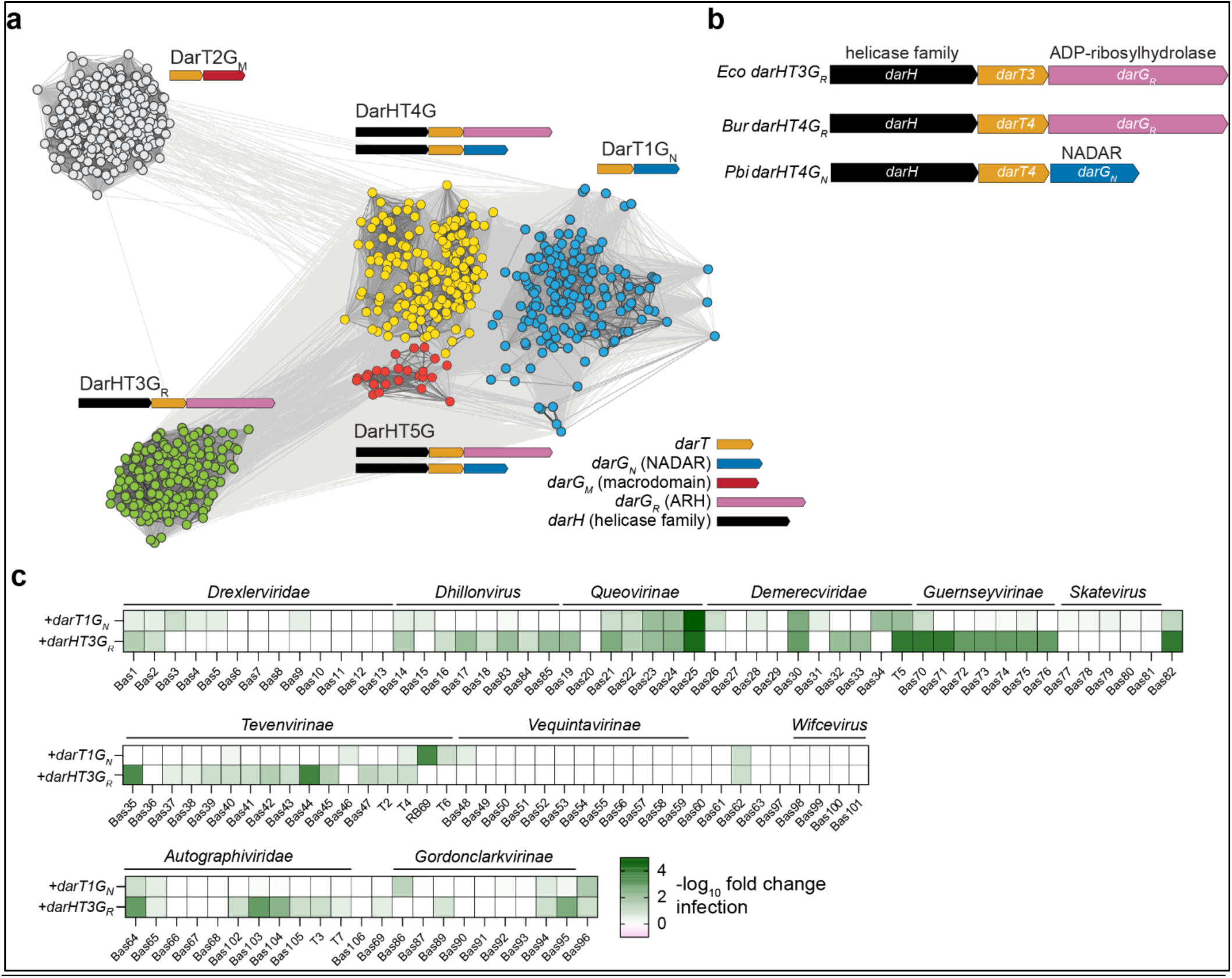
Network clustering analysis reveals DarH-containing DarTG phage defense systems. **a,** Sequence similarity network analysis of DarT proteins. Dots represent proteins, lines represent pairwise sequence similarity. Dots are colored by DarT group. Cartoons below group name indicate representative operon structures within that group (DarT = orange, DarG_N_ = blue, DarG_M_ = red, DarG_R_= pink, DarH = black**. b,** Schematic of DarTG operons selected for further study. Genes colored as in (**a**). **c**, Heatmap showing *darT1G_N_* or *darHT3G_R_* phage defense relative to a strain carrying an empty vector. Darker green indicates stronger defense. Data represent median of 2-6 independent replicates.

We selected three DarHTG systems to study: EcoDarHT3G_R_ (from *E. coli)*, BurDarHT4G_R_ (from *Burkholderia sp. Ax-1719)*, and PbiDarHT4G_N_ (from *Paraclostridium bifermentans)* (Fig. 1b). No group 5 system was selected because this group is small, and clusters closely with group 4. Across all three selected DarHTG systems, DarH has predicted structural similarity with the 1A and 2A RecA motor domains of UvrD, but lacks the regulatory 2B subdomain and the pin motif, elements known to play important roles in DNA unwinding activity (Extended Data Fig. 3)^17–19^. EcoDarT3 has predicted structural similarity to DarT1 and DarT2, and a catalytic glutamate (E167) conserved among all DarTs (Extended Data Fig. 2b,c)^4^. EcoDarG_R_ has a predicted catalytic domain similar to the human ARH1 and ARH3 proteins^20^, with a C-terminal disordered linker and a helix-hairpin-helix domain (Extended Data Fig. 2d).

We first tested whether DarHTG systems have antiphage activity. BurDarHT4G_R_ and PbiDarHT4G_N_ were toxic when induced with a P*_ara_* promoter in *E. coli*, likely due to poor expression of the DarGs which may depend on an internal promoter not recognized in *E. coli*. Thus, we proceeded with EcoDarHT3G_R_, which we cloned under the control of its native promoter and expressed in *E. coli* MG1655. We included EcoDarT1G_N_ (formerly DarTG1) as a comparison. We tested defense against the diverse double-stranded DNA (dsDNA) phages of the Basel collection^21,22^, and additional well-studied dsDNA phages (Fig. 1c). EcoDarHT3G_R_ protected against 43% of the tested phages, with up to a five-log reduction in phage plaquing. In contrast, EcoDarT1G_N_ protected against 16% of the panel; thus, these systems differ in the range and specificity of phage defense.

### DarH enhances DarT activity *in vivo* and *in vitro*

Helicase family proteins are found in many phage defense systems, and often play a role as a sensor or effector protein. To evaluate the necessity of DarH in phage defense, we created an *EcodarHT3G_R_* construct lacking *darH* (*darT3G_R_*) and found that it no longer protected *E. coli* from infection by two representative phages (Fig. 2a). Substituting the conserved catalytic glutamate in DarT for alanine (*darT3^E167A^*) fully abrogated phage defense, indicating that both DarT3 and DarH are necessary for antiviral activity. To distinguish between a sensor or effector role, we tested whether DarH impacts the cellular toxicity of DarT3 in the absence of phage infection. We cloned *EcodarT3* or *EcodarHT3* under the control of an arabinose-inducible promoter (P*_BAD_*) and a theophylline-responsive riboswitch (RS*_theo_*) to enable tighter regulation of expression^23,24^. In *E. coli* BW27783, an MG1655 derivative with a constitutively expressed arabinose importer enabling titratable induction, we found that cell growth was not affected when DarT3 was expressed on its own, but colony forming units (CFU) were reduced by five logs when DarT3 was co-expressed with DarH (Fig. 2b)^25^. While cloning these constructs, we noted that DarT3 inhibited growth in *E. coli* Dh5α cells, which lack the DNA damage repair protein RecA. Notably, deletion of *recA* has previously been reported to sensitize cells to DarT2 toxicity due to impaired ability to repair modified bases^26^. We verified that Dh5α sensitivity to DarT3 was due to the absence of RecA by recapitulating this phenotype in the MG1655-derivative strain BW25113 carrying a *recA* deletion (Fig. 2c)^27^. These data suggest that DarT3 alone is a weakly toxic DNA ADP-ribosyltransferase, and that its toxicity is amplified by co-expression with DarH. To determine if DarT potentiation by DarH is a general feature of helicase-associated DarT systems, we also measured toxicity for BurDarT4 and PbiDarT4 alone or with their cognate DarH and observed a similar dynamic (Extended Data Fig. 4).

**Figure 2:**
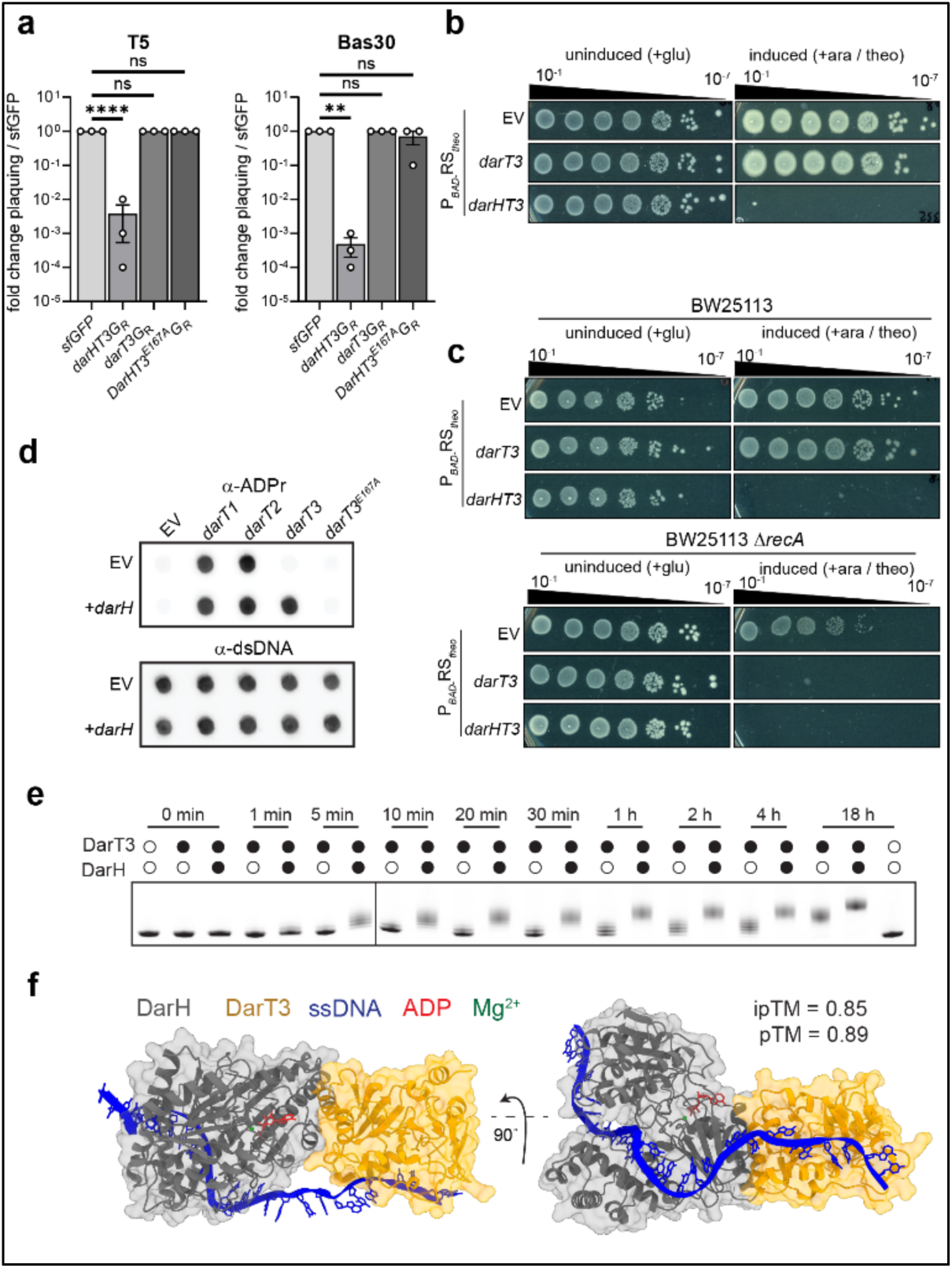
EcoDarH promotes toxin activity in DarHTG systems. **a**, Quantification of phage infection relative to an sfGFP control. Bars represent mean of three experiments, error bars denote standard error of the mean (SEM). Statistical testing was performed by one-way ANOVA with Dunnett’s multiple comparisons test. ****adj. p<0.0001, **adj. p=0.0039. **b**, Representative images showing serial dilutions of *E. coli* plated on glucose (left) or 2mM theophylline plus 0.2% arabinose plates (right). n = 3. **c**, Similar to (**b**), except with *E. coli* strain BW25113 (top) or BW25113Δ *recA* (bottom). **d,** Immuno-dot blot of genomic DNA following expression of the indicated genes. Duplicate membranes were probed with α-ADP-ribose antibody (top) or α-dsDNA antibody (bottom). n = 3. **e,** Representative urea-PAGE gel shift assay showing modification of fluorescein-labeled DNA oligo over time. Open circles indicate absence, filled circles indicate presence of EcoDarH (0.5μM) / DarT (1μM). n = 2. **f**, Predicted structure of EcoDarH (gray) with EcoDarT3 (gold) in the presence of ssDNA (blue), ADP (red), and Mg^2+^ (green). Model on the right is the same, rotated 90 degrees around the X axis.

We then investigated whether EcoDarT3 ADP-ribosylates DNA like other DarTs, and whether this also requires EcoDarH. We measured DarT3 or DarHT3 *in vivo* ADP-ribosylation of genomic DNA (gDNA) by immuno-dot blot, alongside DarT1 and DarT2 controls. DarT3 ADP-ribosylation of DNA was similar to DarT1 and DarT2 when DarH was present, but undetectable without it; this required the catalytic activity of DarT3 (Fig. 2d). To better understand the nature of the potentiation by DarH, we examined the activity of the purified proteins *in vitro* via a time-course ADP-ribosylation gel shift experiment with a ssDNA target (Table 1). DarHT3 rapidly modified the ssDNA, with the first shifted band appearing as soon as one minute post initiation, and all substrate displaying at least one ADP-ribosylation modification by five minutes (Fig. 2e). In contrast, DarT3 alone required four hours to reach this level of modification, and remained below the level of DarHT3 even after 18 hours. We conclude that DarH and DarT3 form a two-part toxin, wherein DarH promotes ssDNA modification by DarT3. AlphaFold3 (AF3) structural modeling predicted a high-confidence interaction (ipTM = 0.85) between DarH and DarT3 in the presence of ssDNA, Mg^2+^, and ADP (Fig. 2f), supporting the idea that these proteins function as a complex to enable efficient DNA ADP-ribosylation.

**Table 1:**
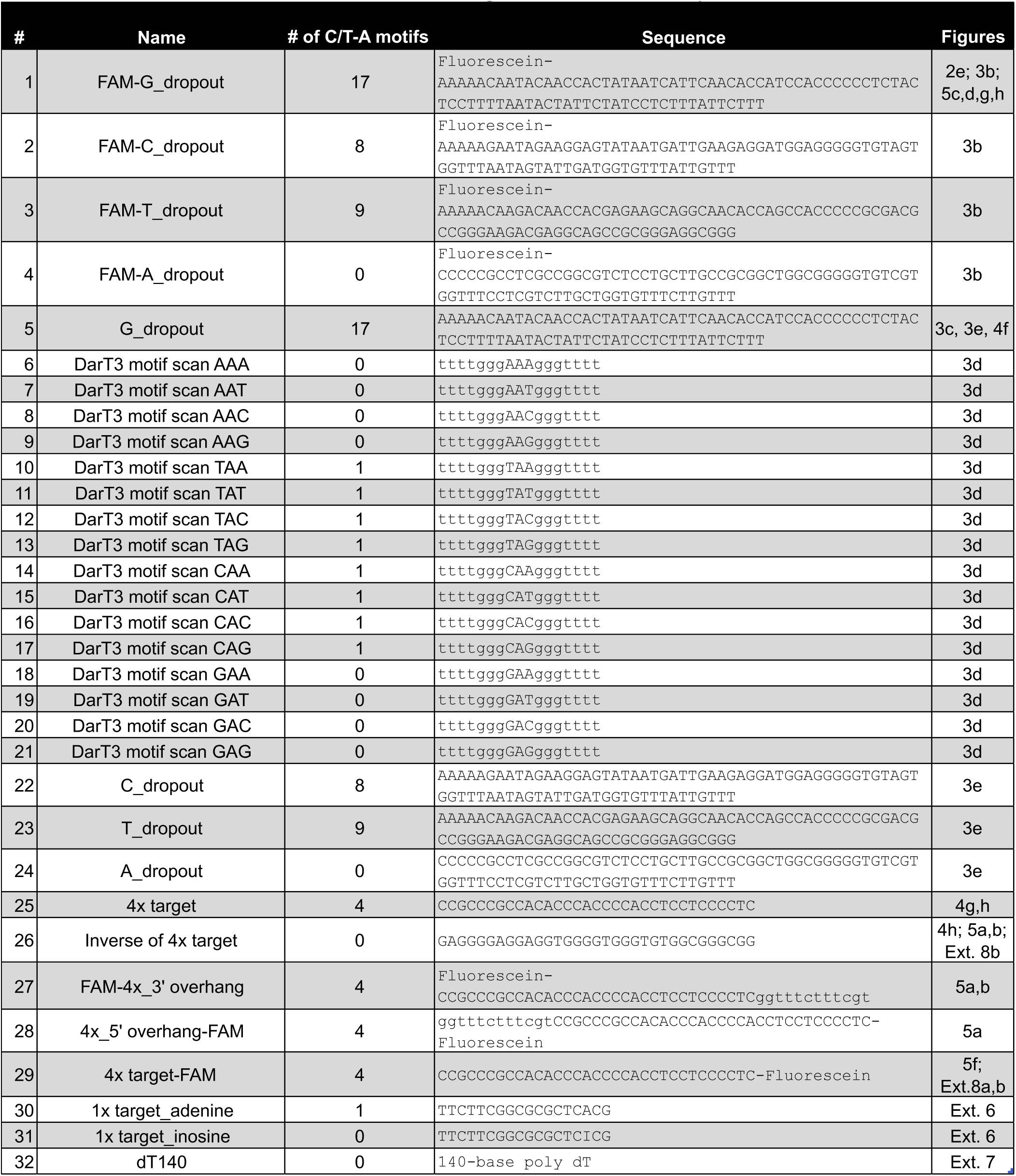
DNA targets used in this study.

### DarHTG systems target DNA adenine or guanine

A defining feature of the *darT1G_N_* and *darT2G_M_* systems is that they target different DNA bases, and therefore their antitoxins are only compatible with their cognate toxins. Because DarT3 clusters apart from DarT1 and DarT2, we tested whether EcoDarG_N_, EcoDarG_M_, or EcoDarG_R_ could neutralize EcoDarT1, EcoDarT2, or EcoDarHT3. As expected, robust bacterial growth was evident when each toxin was expressed with its cognate antitoxin (Fig. 3a). However, neither DarG_N_ nor DarG_M_ was able to rescue growth when co-expressed with DarHT3, implying that DarHT3 targets a distinct DNA base. DarG_R_ partially reversed DarT1 toxicity, as faint colonies grew. Decreasing DarT1 induction resulted in full rescue (Extended Data Fig. 5), suggesting that DarG_R_ can reverse both the major DarT3 target as well as guanine ADP-ribosylated by DarT1, albeit with reduced activity.

**Figure 3:**
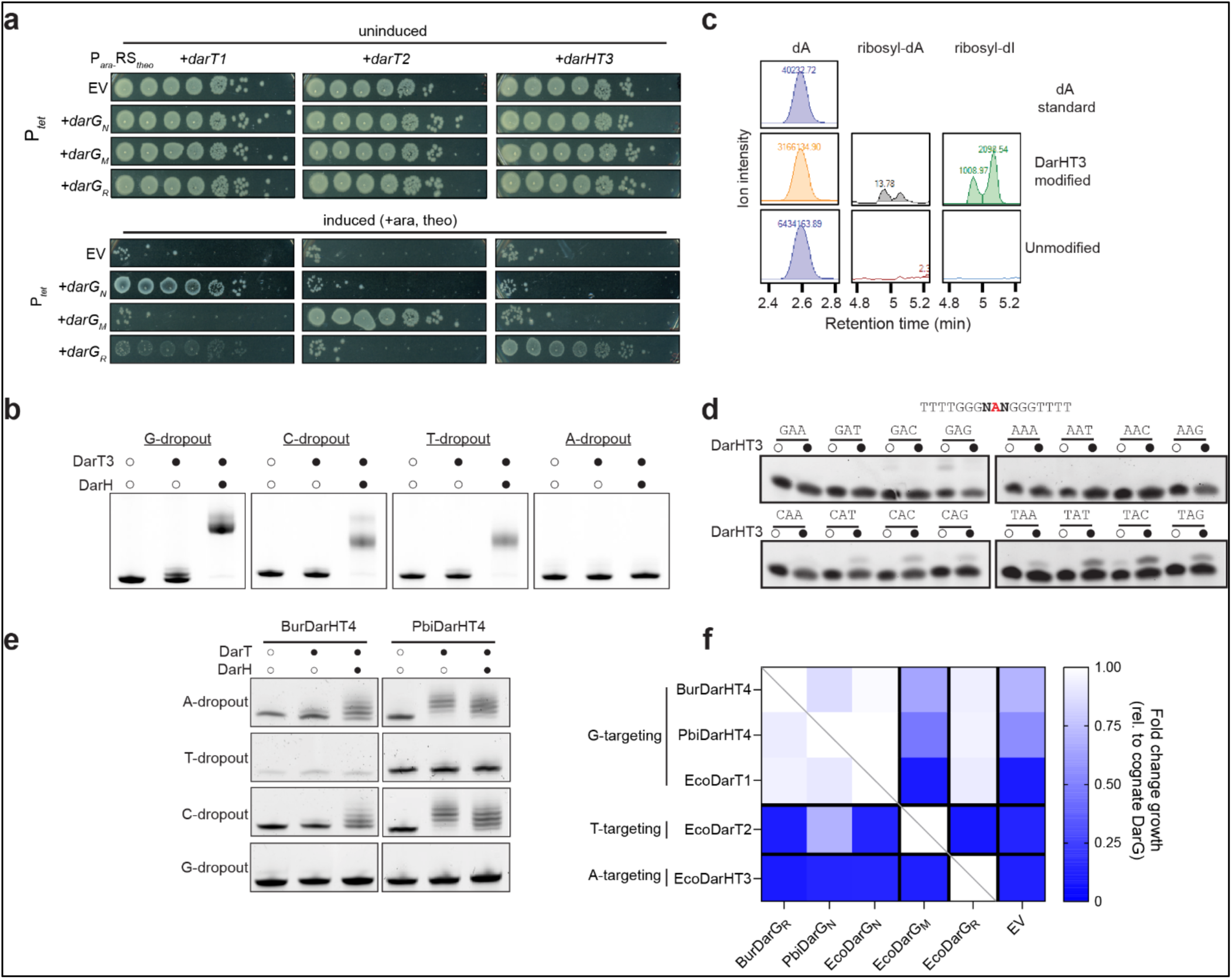
DarHT systems ADP-ribosylate adenine or guanine. **a,** Representative images showing serial dilutions of *E. coli* plated on LB agar with glucose + 10ng / mL anhydrous tetracycline (aTc) or 0.002% arabinose, 2mM theophylline, and 10ng / mL aTc. n = 3. **b,** Representative urea-PAGE gel shift assay with fluorescein-labeled oligos lacking indicated base (“dropout”). Open circles indicate absence, filled circles indicate presence of EcoDarH (0.5μM) / DarT (1μM). n = 3. **c**, HPLC-MS/MS peaks representing retention time (x-axis) and ion intensity (y-axis). Mass transitions were assigned as: 385/137 (for ribosyl-dI), 384/137 (for ribosyl-dA), 252.11/136 (for dA). Representative of two experiments. **d**, Urea-PAGE gel shift assay showing SYBR-gold labeled oligos with a 3-base variable region (in bold). N signifies any nucleotide. Circles indicate presence / absence of DarHT3, as in (**b)**. n = 3. **e,** Urea-PAGE ADP-ribosylation assay showing modification of nucleotide dropout oligos incubated with in vitro synthesized BurDarT4 / HT4 or PbiDarT4 / HT4. **f,** heatmap representing growth of *E. coli* expressing the indicated toxin (rows) and antitoxin (columns). Mean area under the curve was calculated with OD_600_ growth curves, and for each toxin, normalized to the cognate antitoxin (diagonal gray line). White indicates equal growth to cognate antitoxin, blue indicates less growth. Data represent median of three independent replicates.

To identify the target of EcoDarHT3, we performed a gel shift assay with purified EcoDarT3 and four substrate oligos, each lacking one of the DNA bases (Table 1). Every oligo except the one lacking adenine displayed a distinct shift upwards when treated with DarH and DarT3 (Fig. 3b), indicating that the target of group 3 DarTs is likely adenine. To validate this result, we employed high performance liquid chromatography coupled mass spectrometry (HPLC-MS). We modified a ssDNA oligo with DarHT3, then digested it to nucleosides and quantified levels of adenine and ribosyl-adenine relative to an unmodified oligo or adenine standard. We observed the appearance of a ribosyl-A peak after DarHT3 treatment, and a larger peak corresponding to ribosyl-inosine (Fig. 3c). This aligns with a previous report that ADP-ribosylated adenine spontaneously deaminates into ribosyl-inosine in similar workflows and is not biologically relevant^28^. Indeed, when we incubated DarHT3 with an adenine or inosine target oligo, we only observed modification of the adenine target (Extended Data Fig. 6).

To assess whether EcoDarHT3 targets a specific pattern of DNA bases around adenine, we performed a gel shift assay using sixteen oligos which differed only in a three base target site containing every variation of N-A-N, where N is any nucleotide (Table 1). Only targets with a pyrimidine (cytosine or thymine) before the adenine were modified, indicating that the required motif is pyrimidine-adenine (Fig. 3d). The level of modification was highest when the base following the adenine was also a pyrimidine, indicating that EcoDarHT3 prefers a pyrimidine-adenine-pyrimidine motif.

Our clustering analysis indicated that DarT4s are closer to DarT1 (guanine targeting) than DarT3 (Fig. 1a). This group contain a subset of systems with NADAR antitoxins, which have primarily been shown to reverse guanine ADP-ribosylation^2^. We therefore investigated if DarT4 systems target guanine. We generated BurDarHT4 and PbiDarHT4 using *in vitro* transcription and translation (TXTL) followed by in-gel protein detection with Connectase^29,30^ and then performed gel shift assays with our dropout oligo set. Both PbiDarHT4 and BurDarHT4 failed to modify oligos lacking guanine or thymine (Fig. 3e). We inferred that these systems likely modify guanine in a thymine-containing motif. Indeed, growth curve experiments showed that PbiDarG_N_ and BurDarG_R_ were able to prevent EcoDarT1 toxicity, but did not rescue cells from EcoDarT2 or EcoDarT3 (Fig. 3f). Similarly, EcoDarG_N_ and EcoDarG_R_, but not EcoDarG_M_, were able to rescue cells from PbiDarHT4 or BurDarHT4 toxicity. Collectively, these data reveal that DarHT systems target either guanine or adenine, and are found with both ARH and NADAR DarGs.

### EcoDarH enhances EcoDarT3 activity through increased DNA binding and ATP hydrolysis

To determine how EcoDarH enhances EcoDarT3, we considered the core activities of typical helicase family proteins: nucleic acid binding, translocation, and duplex unwinding, with the latter two powered by ATP hydrolysis^19,31^. When we mutated DarH amino acids homologous to those implicated in ATP or DNA binding in UvrD-class helicases^17,32–34^, phage defense was ablated (Fig. 4a-e). These results, consistent with a prior study^16^, suggest that these activities are important for DarH function *in vivo*.

**Figure 4:**
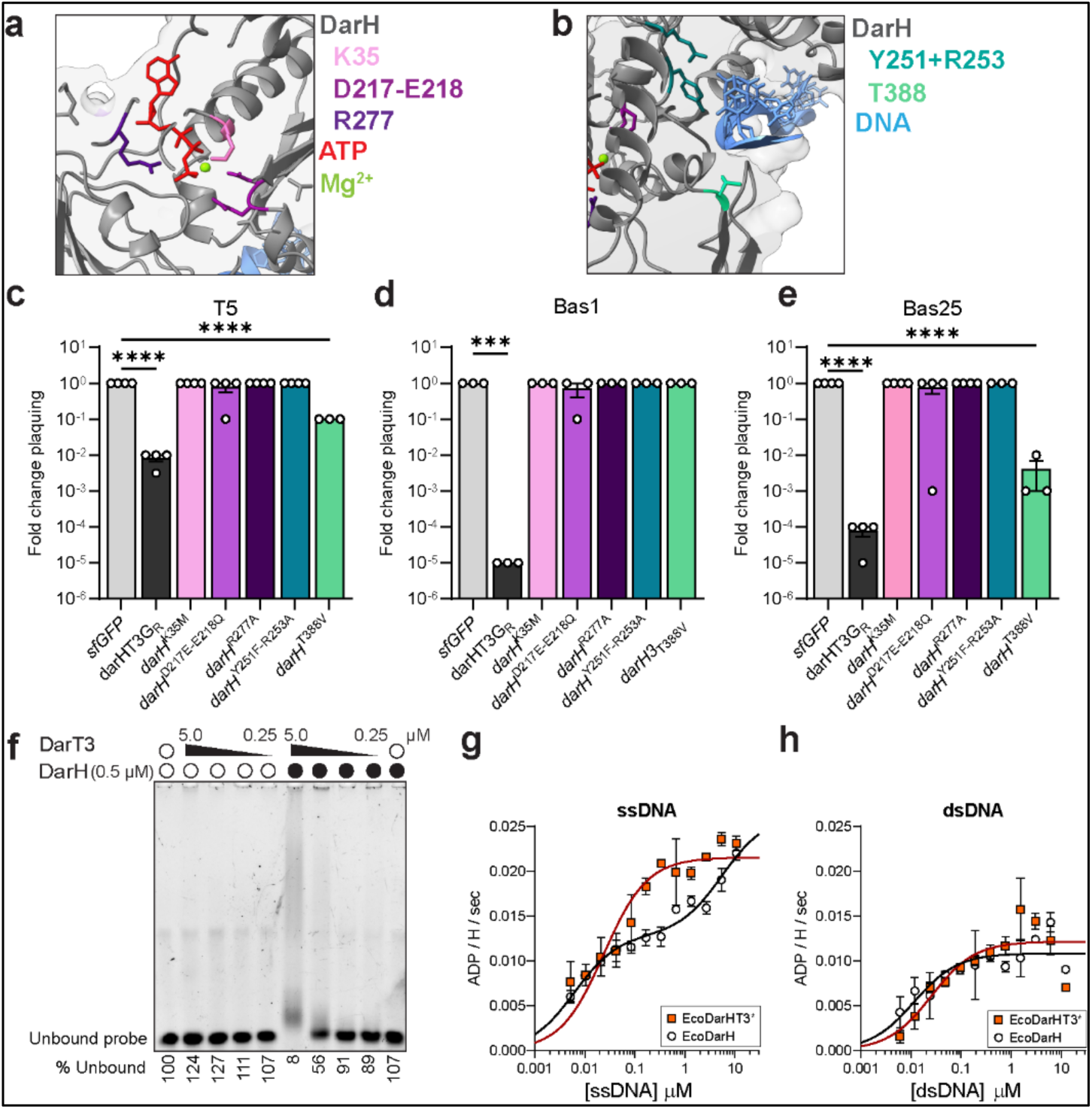
Residues in the DarH ATP and DNA binding sites are required for phage defense. **a,** Cut-away structural model of the EcoDarH (gray) ATP binding site. ATP is shown in red, residue K35 in pink, D217 and E218 in violet, and R277 in purple, and Mg^2+^ in green. **b,** Cut-away model of the EcoDarH DNA binding site. DNA is shown in blue, residues Y251 and R253 in teal, and T388 in green. **c**, Quantification of T5 plaquing (relative to *sfGFP* control). Cells plated on 0.2% arabinose to induce P*_ara_*. Bars represent mean, error bars show SEM of 3-4 replicates. Statistical analysis was performed by one-way ANOVA with Dunnett’s multiple comparisons test. ****adj. p<0.0001. **d**, Quantification of Bas1 plaquing, as in (**c**). ***adj. p=0.0001. **e**, Quantification of Bas25 plaquing, as in (**c**). ****adj. p<0.0001. **f,** EMSA showing EcoDarH / DarT3 interaction with ssDNA. DarT3 was added at 5, 1, 0.5, or 0.25μM, plus or minus DarH at 0.5μM. The percentage of unbound probe normalized to no protein control is shown below the gel. n=3. **g**,**h**, Quantification of ATP hydrolysis by EcoDarH (white circles) or EcoDarHT3^E167A^ (orange squares) in the presence of ssDNA (**g**) or dsDNA (**h**). Points represent mean of two technical replicates, error bars = SD.

To better elucidate the mechanism of potentiation, we examined the DNA binding and ATP hydrolysis activity of DarH *in vitro*. Neither EcoDarT3 nor EcoDarH alone bound stably to a ssDNA probe in an electrophoretic mobility shift assay (EMSA). However, when the two proteins were combined, 92% of the ssDNA shifted upwards, indicating that DarT3 and DarH bind to DNA as a complex, as predicted by AF3 (Fig. 1f, 4f). We next investigated DarH ATPase activity, as predicted ATP-binding site mutants ablated phage defense *in vivo* (Fig. 4c-e). Using an NADH-coupled ATPase assay to follow hydrolysis over time, we found that DarH or DarHT3^E167A^ hydrolyzed ATP in a ssDNA-dependent manner, both achieving a maximum rate of 0.025 ADP molecules produced per DarH per second with 20 μM ssDNA (Fig. 4g). We note that this rate is ∼750-fold slower than the *E. coli* ssDNA translocase and dsDNA helicase UvrD1 (Extended Data Fig. 7). We also tested whether EcoDarH ATPase activity was stimulated by dsDNA, and found a dose-dependent effect, though the maximum rate was half of that achieved with ssDNA, indicating that ssDNA is the preferred substrate (Fig. 4h).

Having shown that the predicted ATP binding site of EcoDarH is required for phage defense *in vivo*, and that DarH displays ATPase activity *in vitro,* we investigated the effect of ATP hydrolysis. EcoDarH lacks structural elements used by UvrD for DNA unwinding (Extended Data Fig. 3), but we tested the possibility that DarH unwinds DNA to generate DarT3 substrate. We designed fluorescein-labeled targets which contained four potential EcoDarT3 modification sites, along with unlabeled complementary strands without pyrimidine-adenine motifs (Table 1). Annealing these oligos generated dsDNA, masking the modification sites unless the duplex was unwound. These targets were blunt-ended, or had 12-base ssDNA overhangs on the 5’ or 3’ side of the target oligo. EcoDarT3 alone did not modify any of the duplex products, but retained its activity on the ssDNA targets (Fig. 5a). The addition of EcoDarH resulted in the appearance of a single faint band above the dsDNA, irrespective of the presence or orientation of a ssDNA overhang. This band was dependent on Mg^2+^, ATP, and NAD^+^, suggesting that it is a *bona fide* product of transient DNA unwinding and ADP-ribosylation (Fig. 5b). However, the low abundance of this band, and the lack of additional bands indicated that DarH is unable to fully or efficiently unwind the dsDNA probes under the tested conditions. It remains unclear whether unwinding of dsDNA is a relevant activity *in vivo*, but in combination with absence of a 2B domain or unwinding pin in DarH and weak binding to dsDNA (Fig. 4h), these data suggest this is unlikely to be its major activity. We conclude that while EcoDarH may enable EcoDarT3 to access a site within dsDNA that DarT3 cannot modify alone, its predominant activity is in accelerating DarT3 ADP-ribosylation of ssDNA.

**Figure 5:**
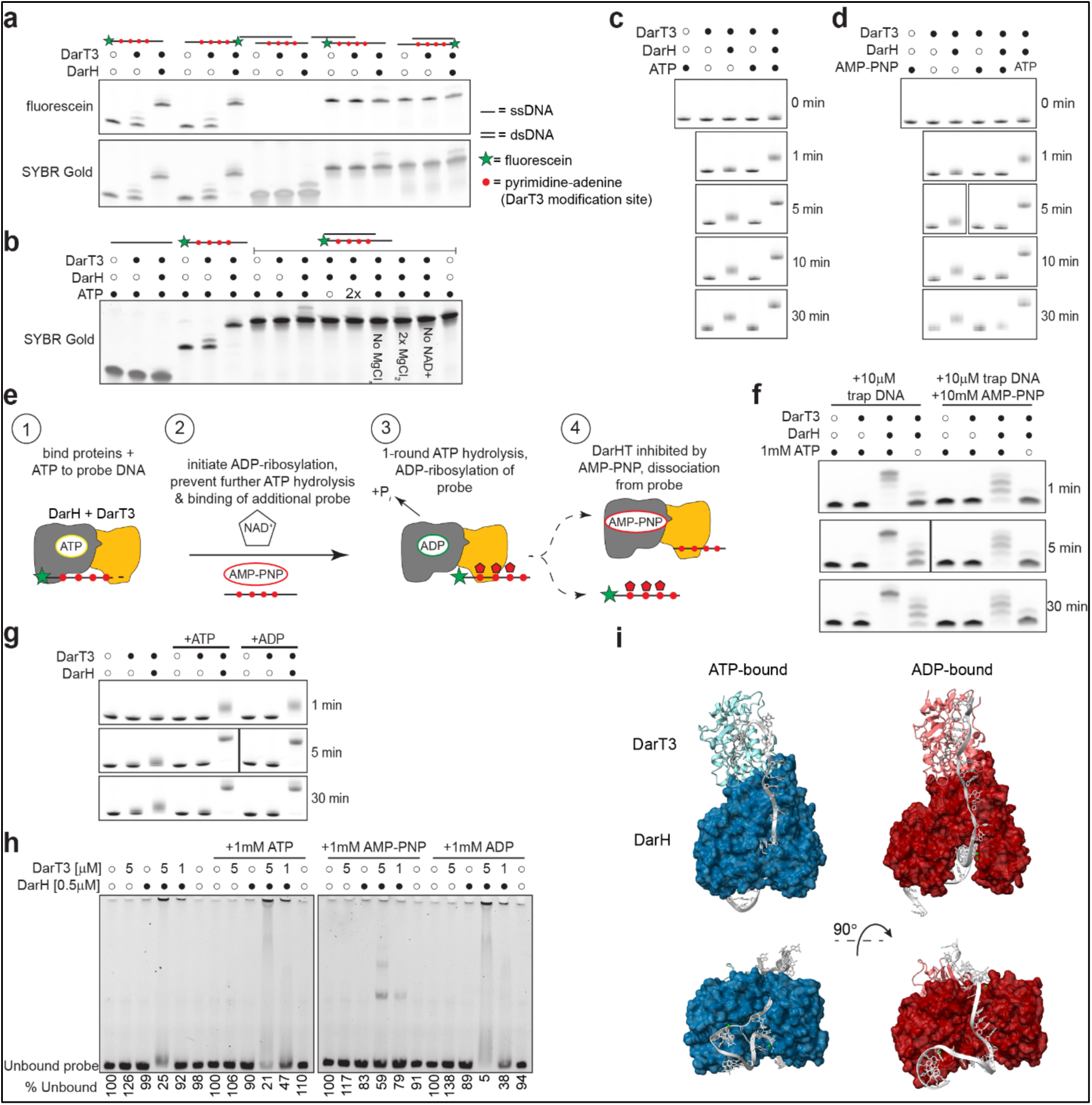
ADP bound EcoDarH promotes EcoDarT3 ADP-ribosylation of ssDNA. **a**, Representative urea-PAGE gel shift assay showing modification of single or double stranded DNA oligos in the presence of 1mM ATP and 1mM MgCl_2_. In schematic, black lines represent DNA strands, red dots indicate potential modification sites, and green stars mark fluorescein labels. Filled black circles indicate presence of EcoDarH (0.5μM) / DarT3 (1μM). n = 3. **b**, similar to (**a**), except with varying levels of ATP, MgCl_2_, and NAD^+^, as marked. **c,d**, Time-course urea-PAGE gel shift assay, showing EcoDarT3 or EcoDarHT3 ADP-ribosylation of ssDNA in the presence or absence of ATP (**c**), or AMP-PNP or ATP (**d**), as indicated by filled circles. n = 2. **e,** Schematic of single ATP turnover ADP-ribosylation assay. Red circles mark potential modification sites, red pentagons represent ADP-ribosylation, and green stars indicate fluorescein. Gray cartoon represents DarH, gold is DarT3. **f,** Time-course urea-PAGE gel shift of fluorescein-labeled DNA after single ATP turnover assay. Presence of EcoDarT3 (1μM), EcoDarH (0.5μM), and ATP are indicated by filled circles. n = 2. **g,** Similar to (**c**), but with ATP or ADP. n = 2. **h,** EMSA showing EcoDarT3 (5 or 1μM, as marked) plus or minus EcoDarH (0.5μM) interactions with 84-base ssDNA in the presence or absence of ATP, AMP-PNP, or ADP. The percentage of unbound probe (relative to no protein control) is displayed below gel. n = 2. **i,** Predicted structures of DarHT3 bound to ssDNA (gray) and ATP (left, in blue) or ADP (right, in red). Green spheres represent Mg^2+^ ions, DarT3-bound NAD^+^ is in dark gray. DarH is shown as a space-filling model. Below is a 90° rotation around the X axis.

As we did not observe evidence of efficient dsDNA unwinding powered by ATP hydrolysis, we tested whether ATP hydrolysis alters EcoDarHT3 ADP-ribosylation of ssDNA. In our initial *in vitro* ADP-ribosylation gel shifts, we saw modification without the addition of exogenous ATP (Fig. 2e), but we reasoned that residual ATP may have been retained by our purified proteins. Therefore, we tested the effect of adding additional ATP and MgCl_2_ to DarT3 or DarHT3. One minute after initiating the reaction, we observed a clear enhancement in DarHT3 activity with ATP, though DarT3 alone was not affected, demonstrating that this increase was dependent on DarH (Fig. 5c). Adding the non-hydrolysable ATP analog adenylyl-imidodiphosphate (AMP-PNP) instead of ATP inhibited the enhancing activity of DarH, suggesting that it is ATP hydrolysis and not ATP binding that activates DarH (Fig. 5d).

UvrD translocates one base per ATP hydrolysis event^31^. To determine whether DarH translocation is coupled to ADP-ribosylation in a 1:1 fashion, we set out to measure the number of ADP-ribosylation events corresponding to one round of ATP hydrolysis. We established a single ATP turnover ADP-ribosylation assay, in which EcoDarH and DarT3 were pre-bound to a fluorescein-labeled probe carrying 4 target sites, ATP, and MgCl_2_, but lacking NAD^+^, the substrate for ADP-ribosylation (Fig. 5e). To initiate ADP-ribosylation, we added NAD^+^; to limit the reaction to a single round of ATP hydrolysis, we also added a 10-fold excess of AMP-PMP to inhibit DarH upon hydrolysis of the preloaded ATP. We also included a 10-fold excess of unlabeled target oligo as a trap to capture DarHT3 complexes upon release from the labeled DNA. Together, these additives allowed us to assess the number of ADP-ribosylation events tied to one cycle of ATP hydrolysis. While trap DNA alone failed to slow down DarHT3 activity, with nearly saturating levels of substrate modification after 1 minute, the addition of AMP-PNP led to laddering, indicating that each probe had been modified between one and four times (Fig. 5f). Furthermore, this pattern was stable out to 30 minutes, suggesting that the reaction was quenched by the additives. We conclude that for every round of ATP hydrolysis, DarHT3 can ADP-ribosylate multiple target sites, leading us to hypothesize that ADP-bound DarH is the active form which enables processive ADP-ribosylation.

To test this, we measured ADP-ribosylation by EcoDarHT3 in the presence of exogenous ADP. ADP enhanced DarHT3 activity as well as ATP, suggesting that the energy of ATP hydrolysis is not required (Fig. 5g). We then investigated whether ATP hydrolysis impacts DarHT3 DNA binding. Adding ATP or ADP with DarHT3 in a ssDNA EMSA increased the amount of probe binding, with smearing observed as in Fig. 4f (Fig. 5h). When we added AMP-PNP, much less DNA was bound, and we observed three distinct bands, suggesting that DarH binds DNA poorly in its ATP-bound state. We observed similar results with a smaller probe, but saw no clear bands with blunt-ended dsDNA (Table 1, Extended Data Fig. 8). In agreement with these experiments, structural modeling of ATP or ADP-bound DarHT3 demonstrate that ADP binding triggers a conformational shift which opens the DarH DNA binding cleft (Fig. 5i).

In sum, our data support a model in which ADP-bound DarH readily binds to ssDNA along with DarT3 (Fig. 6). In this model, ATP hydrolysis does not power directional translocation or unwinding as for a canonical helicase, but functions as a regulatory switch to control effector toxicity.

**Figure 6:**
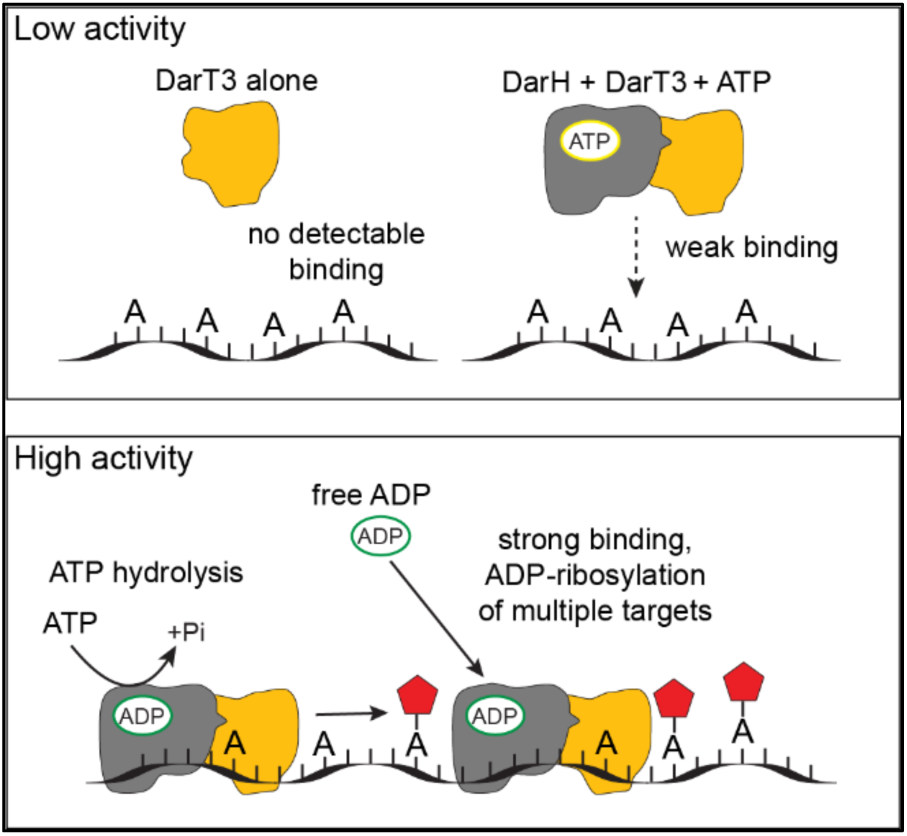
Model of mechanism by which ADP-bound DarH promotes DarT3 activity. Low activity: DarT3 (gold) or ATP-bound DarH (gray) in complex with DarT3 display no or weak binding to ssDNA. High activity: ADP-bound DarH (acquired either via ATP hydrolysis or direct ADP binding) in complex with DarT3 tightly binds to ssDNA, promoting the ADP-ribosylation activity of DarT3 and resulting in high modification levels.

## Discussion

Here, we discover distinct subclasses of DarTG that differ from the previously known DarTG1 and DarTG2 systems in the identity of the antitoxin, the base that the systems target, and the presence of a helicase family protein in the operon. While DarT1 targets guanine and DarT2 targets thymine, we demonstrate that DarT3 targets adenine. Further, DarTs associated with a DarH require it for *in vivo* activity. Under the conditions tested, DarH does not exhibit classical helicase activity, but rather functions to facilitate DarT interactions with ssDNA. This mechanistic insight expands our understanding of the functions that helicase family proteins can exhibit in antiviral defense systems.

Our phylogenetic analysis revealed three new DarH-associated DarT subfamilies, DarT3-DarT5, with DarT3 clustering away from previously studied DarTs. We found that EcoDarHT3 modifies adenine preceded by a pyrimidine. Adenine is a new DarT target, but the CmdT toxin of the CmdTAC defense system ADP-ribosylates adenine in RNA, and human Poly-ADP-ribose Polymerase 1 (PARP1) can mono-ADP-ribosylate adenine in DNA as a minor activity^28,35^. Unlike DarT3, DarT4 and DarT5 cluster close to DarT1, and we found that BurDarT4 and PbiDarT4 modify guanine. We predict that DarT5 toxins also target guanine due to their close association with DarT1 and DarT4. Further, as we found *darG_R_* genes in all DarHTG groups, it is apparent that DarG type does not always correlate with target base, as was previously believed. This is also the first demonstration that ARH family proteins can reverse DNA ADP-ribosylation^20,36^. Our data could enable the development of biochemical and biotechnological tools using adenine targeting enzymes or ARH-class DNA ribosylglycohydolases^37^.

We tested the phage defense activity of *EcodarHTG3* against 116 phages, expanding on the previously published screen of a DarHTG3 system from *Kluyvera sp.*^16^. We find that *darHTG3* defends against a broader set of phages than *darTG1*, which may be due to differences in viral sensing or the prevalence of counter-defense elements that resolve adenine or guanine ADP-ribosylation^38^. The breadth of *darHTG3* defense suggests that it is likely activated by an essential protein or process of dsDNA coliphages. As DarT and DarH are toxic when co-expressed in bacterial cells in absence of phage infection, we speculate that the antitoxin DarG is the sensor of infection, although this mechanism remains under investigation.

Helicase family proteins are prevalent in antiphage systems, often as pathogen sensors or effector proteins^6,10^. We show that DarH, which lacks major DNA unwinding activity and exhibits a low level of ATPase activity in the conditions tested, binds to ssDNA and enhances DarT in the presence of ADP. This suggests that DarH hydrolyzes ATP primarily to generate ADP, rather than to power movement or unwinding of DNA. The multiple ADP-ribosylation events we detect for each round of ATP hydrolysis may be due to the DarHT3 heterodimer sliding or diffusing along DNA, or be the result of multimerization of the complex. The high molecular weight smears we detect in the DNA-protein gel shift assays suggest that DarHT3 may be multimerizing *in vitro*; future studies will investigate whether this also occurs *in vivo* and is relevant for promoting ADP-ribosylation. Regardless of its multimerization status, our data indicate that instead of acting as a molecular motor that ratchets along ssDNA powered by ATP hydrolysis, DarH behaves more like a clamp that tethers DarT to the DNA. DarH joins other proteins such as GajB from the Gabija phage defense system and RIG-I as immune defense helicases which lack major unwinding activity^10,11,39^. It is unclear what advantage DarH confers to a DarTG system, but one possibility is that DarH provides a level of regulation to prevent autoimmune DarT toxicity.

Overall, this work details how a helicase family protein enhances a phage defense effector, which may inform studies of the many uncharacterized immune-associated helicases. We also expand the known arsenal of DarTG defenses, including new targets for DNA ADP-ribosylation and enzymes to remove this mark.

## Methods

### Strains and growth conditions

For a complete list of bacterial strains, see Supplementary Tables 1. *E. coli* DH5α cells were used for plasmid construction, while *E. coli* Rosetta2(DE3) cells were used for protein expression, and *E. coli* MG1655, BW27783, or BW25113 were used for all other experiments. The latter two were used for all experiments with arabinose-inducible plasmids. *E. coli* were grown at 37°C in Lysogeny Broth (LB) unless otherwise noted. Plasmids were transformed by the TSS method^40^ or Zymo Mix & Go chemical transformation (Zymo Research #T3001). Transformants were selected on LB agar plates plus carbenicillin (100 μg ml^−1^), kanamycin (50 μg ml^−1^), or spectinomycin (50 μg ml^-1^). In liquid LB cultures, plasmids were maintained with carbenicillin (50 μg ml^−1^), kanamycin (30 μg ml^−1^), or spectinomycin (50 μg ml^-1^). For plasmids with an arabinose promoter, gene expression was suppressed during cloning and propagation with glucose (0.4% w/v). To induce transcription, arabinose (0.2% w/v) was added. For plasmids encoding a theophylline riboswitch, translation was induced with 2mM theophylline in 0.1M NaOH. For plasmids with a Tet promoter, transcription was induced with 10ng/mL aTc. For plasmids with a lac promoter, transcription was induced with 0.1mM IPTG.

Phage stocks were propagated on *E. coli* MG1655 (Bas1-69, T2-T7, RB69), K12 Δ*RM wbbL*+ (Bas70-73, 77-80), or BW25113Δ*gtrS wbbL*+ (Bas74-76, Bas81-106). Host cells were grown to approximately OD_600_ = 0.2-0.3 with shaking, then infected at a Multiplicity of Infection (MOI) of 0.01-0.1. Cultures were shaken at 37°C until clearance or overnight, and remaining cells and debris were pelleted at 4,000 x g for five minutes. Supernatants were filtered through a 0.22μm membrane and stored at 4°C.

### Plasmid construction

To synthesize DarTG variant systems, DNA fragments containing the target gene were ordered from IDT DNA, GenScript Biotech, or Twist Bioscience. Linearized vector and / or subcloned genes were amplified through polymerase chain reaction with KAPA high-fidelity polymerase (Roche Diagnostics #9420398001) using primers containing 25-base overlaps with the target site. PCR products or synthesized gene fragments were inserted into target vectors by Gibson assembly, using NEBuilder HiFi DNA assembly (New England Biolabs #E2621). For mutagenesis, overlapping primers including the mutation(s) were used with corresponding overlapping primers in another region of the plasmid to generate two PCR products with overlapping ends, including the mutations. These products were joined by Gibson assembly. Assembled plasmids were transformed into chemically competent DH5α cells and selected on LB plus the appropriate antibiotic. Plasmids were verified by whole-plasmid sequencing. For phage infection experiments with DarTG expression controlled by the native promoter, defense systems were cloned into the pBR322 vector. The pJB37-P*_ara_* vector was used for expression of complete *darTG* systems controlled by an inducible promoter. The expression of toxins required stricter control of expression, and so *darT* or *darHT* genes were cloned into pJB37-P*_ara_*-Riboswitch_Theophylline_. Non-toxic proteins (darH, darG) were cloned into pKVS45-P*_tet_.* For protein purification, genes were cloned into a pET-P*_lac_* plasmid with a T7 promoter. Lists of primers and gene fragments used in this work are available in Supplementary Table 2. Plasmids were designed and sequences were aligned using SnapGene v8.2.2.

### Sequence searches, sequence similarity network analysis, and gene neighborhood analysis

An initial gene neighborhood analysis of DarT was performed using fast.genomics^13^, which searches homologs in a selected set of prokaryotic genomes and annotates the neighboring genes based on pre-generated sequence clusters using CD-HIT. To systematically study sequence relationship and associations of the DarT homologs, we performed iterative PSI-BLAST searches^14^ against the NCBI non-redundant (nr) protein database using several known DarT proteins as queries. The searches were performed with an e-value cut-off of 0.005 and continued until convergence. To identify the major groups of DarT proteins, we conducted sequence similarity network analysis using CLANS^15^. In this analysis, all-against-all BLASTP comparisons were conducted using the BLOSUM62 substitution matrix and an e-value cutoff of 0.0001. The resulting pairwise similarities were visualized as a two-dimensional network, with nodes representing sequences and edges representing detected pairwise similarities. The network layout was generated using by the Fruchterman and Reingold force-directed graph drawing algorithm^41^. Colors and labels were added using Adobe Illustrator.

To examine the genomic context of the major DarT groups identified by network analysis, we retrieved their four upstream and four downstream gene neighbors from the NCBI GenBank database^42^. Proteins encoded by these neighboring genes were clustered by sequence similarity using the BLASTCLUST program, a BLAST score-based single-linkage clustering method (https://ftp.ncbi.nih.gov/blast/documents/blastclust.html). The resulting protein clusters were annotated based on their domain architectures which were identified using HMMSCAN^43^ searching against the Pfam database^44^ and our in-house HMM profile database. Signal peptides and transmembrane regions were predicted using Phobius^45^.

### Plaque assays

Overnight cultures of *E. coli* were diluted 1:20 in melted top agar (LB + 0.5% agar + inducers, as required) and overlaid on top of LB + 1.2% agar plates. Phages were diluted from 1 × 10^−1^ to 10^−8^ in LB (Basel panel) or sterile PBS (all other plaque assays). Once top agar had solidified, phages were spotted in 2μL droplets and air-dried. Infected plates were incubated overnight at 37°C for most phages, or at 30°C (Bas103). Plaque forming units per mL (PFU/mL) values were calculated by counting plaques, although sometimes plaques were too small to count in the presence of DarTG systems. In these cases, the fold change in infection relative to a control strain was estimated from the last dilution with visible plaques.

### Colony forming unit (CFU) spotting assays

Overnight cultures of *E. coli* were diluted ten-fold from 1 × 10^−1^ to 10^−8^ in LB, and spotted onto LB plates with or without inducers. Plates were air-dried and incubated overnight at 37°C. Colony counts were used to calculate CFU / mL.

### DNA immuno-dot blot

Overnight cultures of E. coli were back-diluted 1:100 into fresh LB plus glucose to suppress toxin expression, and incubated in a shaking incubator at 37°C until reaching an OD_600_ = 0.4. Glucose was washed off and cells were resuspended in fresh LB plus arabinose and theophylline to induce toxin expression, and aTc to induce *darH* expression. Cells were shaken for another 20 minutes, then pelleted and genomic DNA was extracted using a Puregene Cell kit (Qiagen # 158043). dsDNA was quantified using a Qubit fluorometer, and 2.5μg of DNA was fragmented by digestion with 5 units of AluI restriction enzyme as previously described^46^ (NEB #R0137). Samples were spotted onto a nitrocellulose membrane, in duplicate, using a Biorad Bio-Dot vacuum manifold (1μg DNA per well). DNA was crosslinked to the membrane by UV irradiation, and membranes were blocked in Tris-buffered Saline + 0.1% Tween-20 + 5% milk. Duplicate membranes were separated and incubated with primary antibodies overnight, rocking at 4°C. Mono-ADP-ribosylation was detected with Horseradish peroxidase-conjugated anti-MonoADP-ribose Fab (1:10,000, Biorad #TZA020P). Double stranded DNA was detected with mouse anti-dsDNA (1:1000, DSHB #autoanti-dsDNA), followed by secondary labeling with anti-mouse HRP (1:10,000, Invitrogen #32430). After washing with TBST, blots were developed with Immobilon ECL substrate (anti-ADP-ribose, Millipore Sigma #WBKLS0050) or SuperSignal femto max sensitivity HRP substrate (anti-dsDNA, ThermoFisher #34095).

### Protein purification

*E. coli darH* and *darT3* were cloned into separate pET expression vectors with a C-terminal Tobacco Etch Virus (TEV) protease cleavage site followed by a 3x Flag tag and 10x His tag. Overnight cultures were back-diluted 1:100 and grown to an OD_600_ of ∼0.5 at 37°C, and 0.1mM IPTG was added to induce expression. Cells were shaken for another two hours at 30°C, then cells were pelleted and resuspended in lysis buffer (50mM HEPES, 250mM NaCl, 5% glycerol, 20mM imidazole, 50 units/mL Benzonase (Millipore-Sigma E1014), and 1x cOmplete protease inhibitors (Roche #4693132001)). Cells were disrupted by sonication on ice and lysates were clarified by centrifugation at 25,000 x g for 30 minutes, at 4°C. His-tagged proteins were bound to nickel-NTA agarose beads (Goldbio Biotechnology #H-350) in gravity flow columns (Goldbio Biotechnology #P-301) at 4°C, and beads were washed five times with 12 mL of wash buffer (50mM HEPES, 250mM NaCl, 10% glycerol, 20mM imidazole, 0.5% Triton X-100). Bound proteins were eluted in elution buffer plus a stepwise increase in imidazole (1x PBS, 5% glycerol, 100 / 200 / 300 / 400mM imidazole), and fractions were analyzed for protein content by PAGE and Coomassie staining. Fractions with protein at the correct size were dialyzed into storage buffer 1 or 2 overnight (storage buffer 1: PBS, 5% glycerol. Storage buffer 2: 50mM HEPES, 100mM NaCl, 5% glycerol).

### ADP-ribosylation gel shift assay

Purified DarT3 (1 μM) and / or DarH (0.5μM) were mixed with ADP-ribosylation buffer (50mM Tris-HCl [pH 8], 50mM NaCl), 0.5mM NAD^+^, and 3μM DNA probe. For experiments investigating ATP hydrolysis, 1mM MgCl_2_ plus 1mM ATP, ADP, or AMP-PNP was also added. Reactions were incubated at 37°C for 30 minutes (unless otherwise indicated). For time course experiments, reactions were stopped at the indicated time by heating at 95°C for 5 minutes. TBE-Urea sample buffer (Invitrogen LC6876) was added and samples were denatured at 70°C for three minutes before electrophoresis on 7M urea, 15% acrylamide TBE gels at 180V for 90 minutes, at 4°C. For fluorescein-labeled probes, gels were directly imaged on a ChemiDoc system, or for unlabeled probes, DNA was stained with 1x SYBR Gold (Invitrogen #S11494) before imaging.

### High-Performance Liquid Chromatography combined with Mass Spectrometry (HPLC-MS/MS)

DNA oligos incubated with or without DarH and DarT3 were digested with 2 μL of Nucleoside Digestion Mix (NEB, M0649S) in a total volume of 50 μL at 37°C overnight. The digested sample was filtered with a 0.22 μm filter and then processed for nucleoside mass spectrometry analysis^47,48^. Chromatographic separation was performed using an Agilent 1290 Infinity II UHPLC system with a ZORBAX RRHD Eclipse Plus C18 2.1 x 50 mm (1.8 μm) column. The criteria settings for HPLC and the detection of ribosyl-deoxynucleosides by MS were adapted from previous work^28^. The mobile phase consisted of water and methanol (with 0.1% formic acid) run at 0.5 mL/min. The run started with a 3 min gradient of 2-8% methanol, followed by a sharp increase to 98% methanol which was maintained for another 5 min. Mass spectrometric detection was performed using an Agilent 6470 Triple Quadrupole system operating in positive electrospray ionization mode, monitoring the mass transitions 385/137 (for ribosyl-dI), 384/137 (for ribosyl-dA), 252.11/136 (for dA). The data were visualized with the Agilent Qualitative Navigator analysis software.

### ADP-ribosylation assays in cell-free systems

DarT and DarH, each carrying a C-terminal Cn-tag, were expressed separately from linear DNA templates using the myTXTL system (Arbor Biosciences#5410ML) according to the manufacturer’s protocol. Coding sequences were placed under the control of the P70 promoter. Templates were added to the TXTL mixture at a final concentration of 6 nM, and reactions were incubated for 16 h at 29 °C. Mock TXTL reactions were prepared without an expression template.

Protein expression was verified by Connectase-mediated in-gel fluorescence labelling, as described previously^29,30^. Briefly, 1 µL of each cell-free expression reaction was mixed with 1 µL of labelling solution containing 20 nM Connectase and 20 nM Cy5.5 fluoropeptide (RELASKDPGAFDADPLVVEISEEGE-Cy5.5) and incubated for 20 min at 22 °C. Subsequently, 1 µL of 3× Reducing Blue Protein Loading Dye (NEB#B7703) was added. Samples were heated at 95 °C for 5 min and immediately transferred to a frozen cooling rack. The entire sample volume (3 µL) was loaded onto precast Mini-PROTEAN TGX Any kD gels (BioRad#4569035) and electrophoresed for 120 min at 100 V. Gels were imaged using a ChemiDoc MP Imaging System (BioRad#12003154) with Cy5.5-compatible acquisition settings, and images were processed using Image Lab software (v6.1.0). For the *in vitro* ADP-ribosylation assays, we adjusted a protocol by Gupta et al, 2025^37^. Briefly, ADP-ribosylation assays (5 µL) contained 25% (v/v) TXTL mixture, consisting of equal volumes of DarT-expressing TXTL and either DarH-expressing or mock TXTL. Mock controls contained 25% (v/v) mock TXTL only. Final assay concentrations were 12.5 mM Tris–HCl (pH 8.0), 37.5 mM NaCl, 50 µM NAD⁺, 10 mM EDTA and 10 µM of the indicated single-stranded dropout oligonucleotide. Reactions were incubated for 1 h at 30 °C.

Samples were subsequently treated with RNase A (0.5 µg per sample; NEB#T3018L) for 30 min at 37 °C. Proteinase K TL (1 µL; 0.12 U; NEB#P8111S) and nuclease-free water were then added to bring each sample to a final volume of 50 µL, yielding a final Proteinase K TL concentration of 2.4 U/mL. Samples were incubated for 15 min at 25 °C. Oligonucleotides were purified using the Oligo Clean & Concentrator kit (Zymo Research#D4061), diluted and mixed with an equal volume of RNA loading dye 2X (NEB#B0363S). Samples were heated at 95 °C for 5 min and immediately transferred to a frozen cooling rack. Approximately 8 ng of ssDNA per well was loaded onto 20% polyacrylamide gels containing 8 M urea and resolved by denaturing PAGE for 150 min at 150 V. Gels were stained with SYBR Gold (Invitrogen#S11494) according to the manufacturer’s protocol and imaged using the ChemiDoc MP Imaging System with SYBR Gold-compatible acquisition settings.

### Growth curve toxicity assay

Overnight cultures of bacteria were back-diluted 1:100 into fresh LB plus 0.2% arabinose, 2mM theophylline, and 10ng/mL aTc. These dilutions were plated in triplicate (100μL volume) in a propylene, round-bottom 96 well plate (USA Scientific #1830-9610), and overlaid with 70 μL of mineral oil to prevent evaporation. Plates were sealed with a gas-permeable membrane (USA Scientific #9123-6100) and incubated for 8 hours at 37°C with orbital shaking at 432 rpm on a Tecan Infinite M200 Pro plate reader. Optical density was read at 600nm every ten minutes. The mean absorbance from triplicate media-alone wells was subtracted from all other wells to calculate blank-subtracted OD_600_.

### Electrophoretic mobility shift assay

Proteins were diluted in phosphate buffer (50mM sodium phosphate [pH 7.5] plus 200mM NaCl), along with 1mM ATP, ADP, or AMP-PNP and MgCl_2_, as indicated. DNA was added to a final concentration of 50nM in binding buffer (10mM sodium phosphate [pH 7.5], 100mM NaCl, 0.5mM DTT, 40ug/mL BSA), and the reactions were incubated at 30°C for 30 minutes. 5x loading dye (Tris-borate (TB) buffer, 15% Ficoll-400, 0.1% bromophenol blue, 0.1% xylene cyanol) was added and the samples were run on 6% polyacrylamide DNA retardation gels (Invitrogen #EC6365BOX) or hand-cast 6% polyacrylamide Tris-Borate gels, at 100V for 1.5 hours at 4°C. For fluorescein-labeled DNA probes, gels were directly imaged on a ChemiDoc system, or for unlabeled probes, DNA was stained with 1x SYBR Gold (Invitrogen #S11494) before imaging.

### Fluorescence-coupled ATPase assays

Steady-state ATP hydrolysis was measured by coupling the production of ADP to the reduction of NADH via the glycolytic enzymes pyruvate kinase and lactate dehydrogenase in a fluorescent plate reader assay^49^. ATPase reactions with DarH alone or in the presence of DarT3 were carried out in Reaction buffer (50 mM TrisHCl pH 8, 50 mM NaCl, 1 mM MgCl2) at 37°C. The reported concentrations are the final concentrations after mixing. Reactions were prepared in black, 384 low volume, round bottom well plates (Corning #4514). Two-fold serial dilutions of DNA substrates were prepared then mixed with a solution containing 0.133 mg/ml NADH (β-nicotinamide adenine dinucleotide reduced, Sigma #N4505), 5.6 U/ml pyruvate kinase/ 7.8 U/ml lactate dehydrogenase (Sigma #P0294), and DarH (0.5 μM, or with DarT3, 1 μM), and incubated at 25°C for 15 min. The plate was placed in a CLARIOstarPlus fluorescent plate reader (BMG LabTech) and incubated for 5 min at the reaction temperature. After an initial fluorescent intensity mode scan of the plate (excitation: 357nm, emission: 466nm, bandwidth 20nm for both), the reaction was initiated by adding 0.31 mg/ml PEP (phosphoenolpyruvic acid, Sigma #860077) and 0.5 mM ATP using the plate reader injector (430 μL/sec). The plate was then rapidly mixed (10 sec, double orbiter, 600 rpm) and scanned over a period of 30 minutes. The fluorescent signal was calibrated for each plate of reactions by setting up a series of reactions with known amounts of ADP without DarH and DNA. All reactions were prepared in duplicate. After converting the fluorescent signal to moles of ADP, the initial rate was measured by taking the linear portion of the time course within the first 10 minutes.

### Single-ATP turnover ADP-ribosylation assay

Purified DarT3 and/or DarH were mixed with 1μM fluorescein-labeled probe DNA, 1mM MgCl_2,_ and 1mM ATP in ADP-ribosylation buffer. This mix was incubated at 30°C for 10 minutes, then cooled to 4°C. To initiate ADP-ribosylation, a mix of 0.5mM NAD^+^, 10mM AMP-PNP, and 10μM unlabeled target oligo was added. Reactions were incubated at 37°C for the indicated times, and gels were run and imaged as for other ADP-ribosylation gel shifts.

### Structural prediction / display

Protein structures were predicted using AlphaFold server^50^. Outputs were visualized using UCSF ChimeraX version 1.10.1^51^. Structures were aligned and root mean square deviation values were computed using the Matchmaker tool with default settings.

### Statistical analysis

Statistical analyses were performed using GraphPad Prism version 11.0.2. For comparing multiple groups to a control, one-way ANOVA with Dunnett’s multiple comparisons tests were used. For analysis of growth curves, three technical replicates were plotted per sample, and an area under the curve analysis was used to calculate total area.

## Acknowledgements

This work was supported by an NIAID R01 (R01AI195977) and unrestricted funds from Washington University School of Medicine to M.L. T.G. was supported by a NIAID F32 (F32AI202943). C.P., A.Q.A and S.S.K. acknowledge funding from the Research Council of Lithuania under the Programme University Excellence Initiatives of the Ministry of Education, Science and Sports of the Republic of Lithuania (Measure No. 12-001-01-01-01, Improving the Research and Study Environment), project No: S-A-UEI-23-10. C.P. and A.Q.A. further acknowledge funding from the EMBO Installation Grant programme under grant No 6357. Y.T. and D.Z. were supported by an NIAID R21 (R21AI188314) and Saint Louis University President’s Research Fund. We appreciate the support of M. Williams and the Molecular Microbiology Media and Glassware Facility. We also thank Dr. Patrick Pausch for his advice on protein structures. We thank all members of the LeRoux lab for comments and discussions.

## Author Contributions

Conceptualization, M.L., T.G.; Data Curation, D.Z.; Formal Analysis, M.L., T.G., D.Z.; Funding Acquisition, M.L., T.G., C.P., D.Z.; Investigation, T.G., E.T., A.A., Y.T., J.T., S.K., N.C.; Methodology, M.L., T.G., C.P., E.T., E.G., D.Z., N.M., N.T.; Supervision, M.L., C.P., D.Z., N.M., E.G.; Visualization, T.G., D.Z., A.A., M.L.; Writing – original draft, M.L., T.G.; Writing – review & editing, M.L., T.G., J.T., C.P., A.A., E.T., E.G., N.M., N.T., D.Z.

---

**Extended Data Figure 1:**
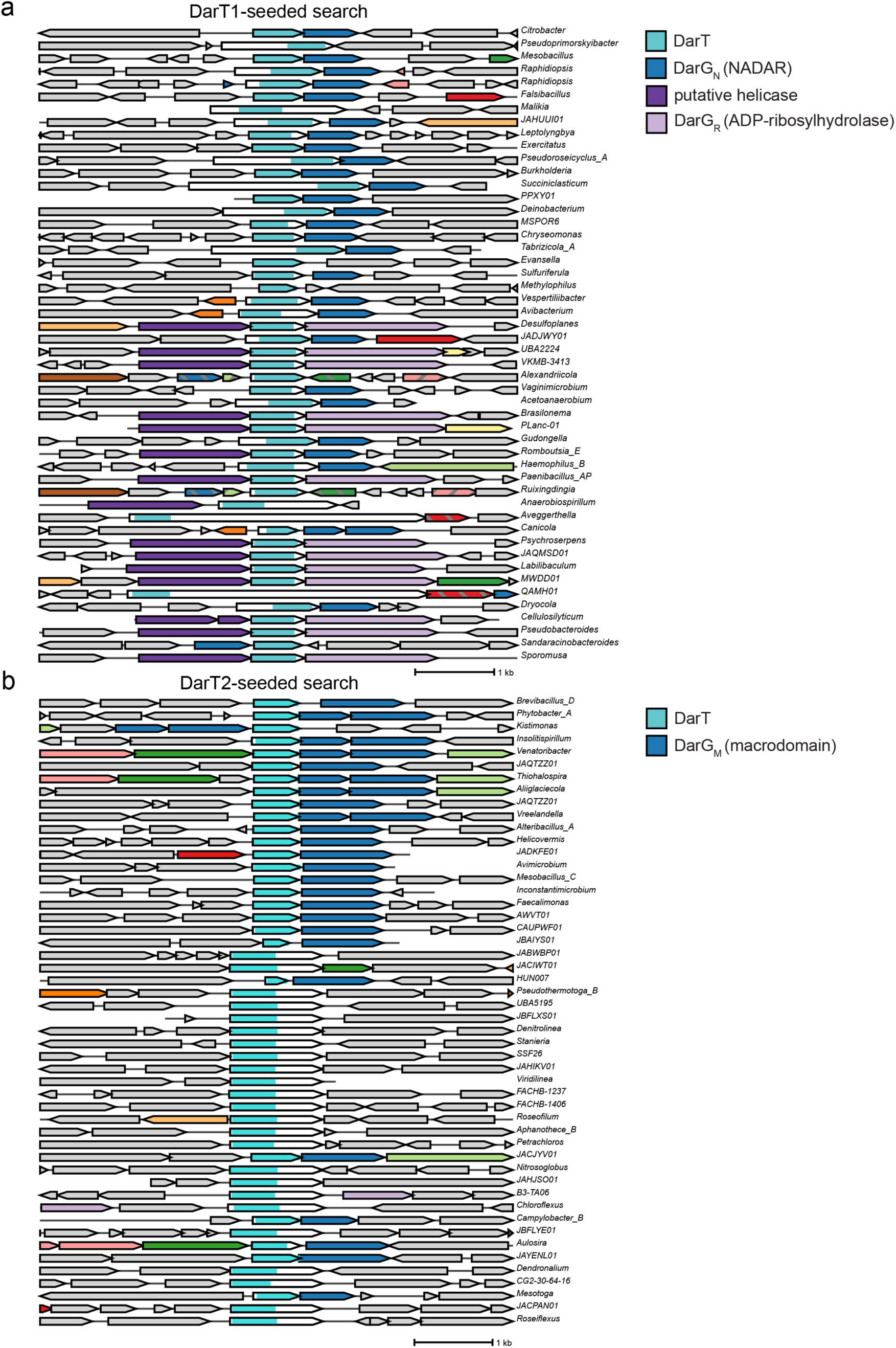
Gene neighborhood analysis reveals existence of DarHTG systems. **a**, Output of a fast.genomics search seeded with the amino acid sequence of *E. coli* DarT1. DarT homologs are shown in sky blue, DarG1 in dark blue, helicase family proteins in purple, and ARH domains in light purple. **b**, Similar to (**a**), but seeded with *E. coli* DarT2. DarT2 homologs are shown in sky blue, DarG2 in dark blue.

**Extended Data Figure 2:**
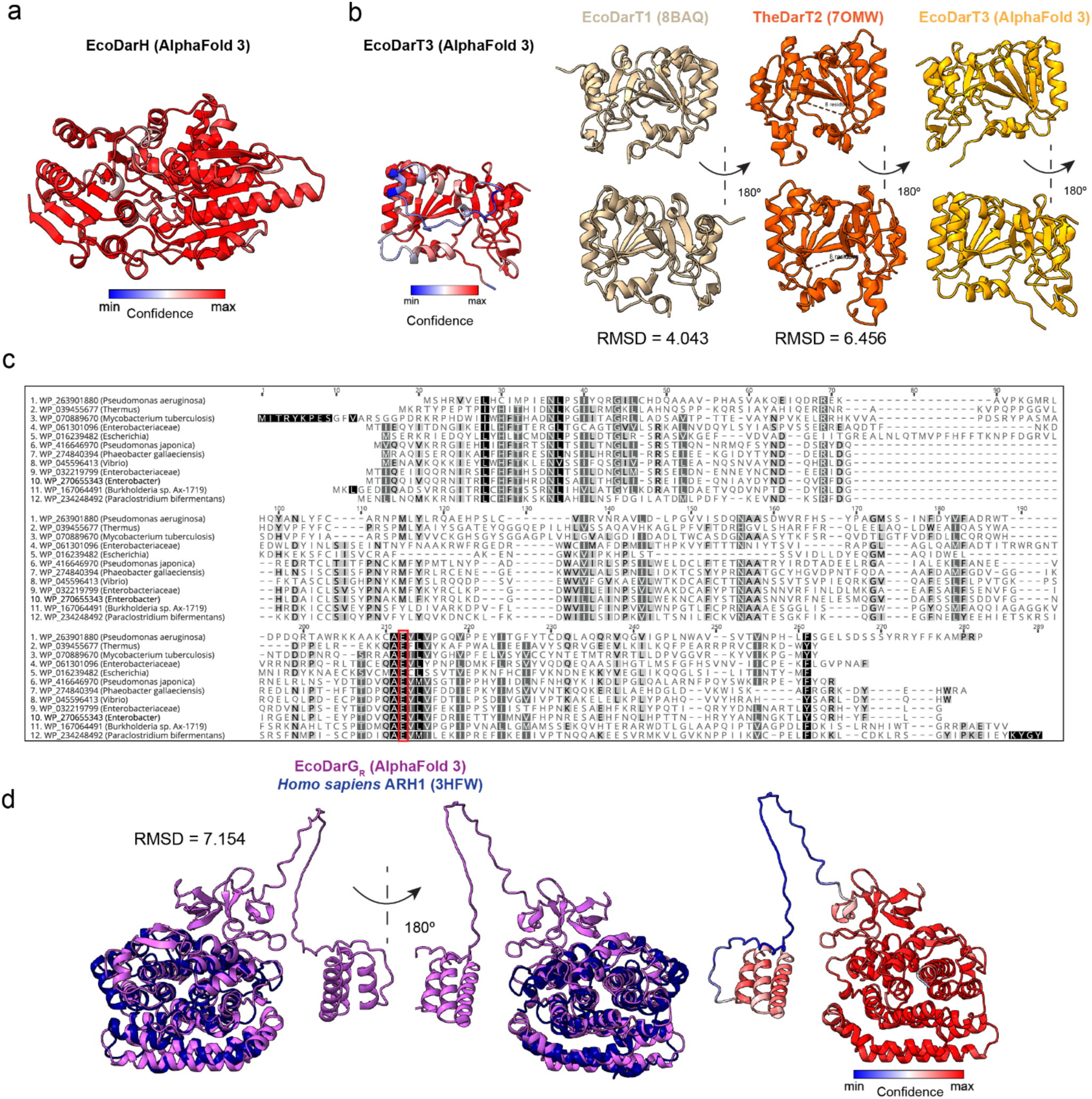
Predicted structures of *E. coli* DarH, DarT3, and DarG_R_. **a,** AlphaFold3 predicted structure of DarH, colored by pLDDT. **b**, Structures of *E. coli* DarT1 (tan), *Thermus sp.* DarT2 (orange), and predicted structure of *E. coli* DarT3 (gold or colored by pLDDT). Bottom row is a 180 degree rotation of top row. **c,** Amino acid sequence alignment of representative DarT1-4 proteins generated by Geneious (v2025.1.3). Columns are colored by % identity, where darker is more conserved. The catalytic glutamate residue is outlined in red. **d,** Predicted structure of *E. coli* DarG_R_ (violet) overlaid with *Homo sapiens* ARH1 (blue). Middle is a 180 degree rotation, while the right is DarG_R_ alone, colored by pLDDT.

**Extended Data Figure 3:**
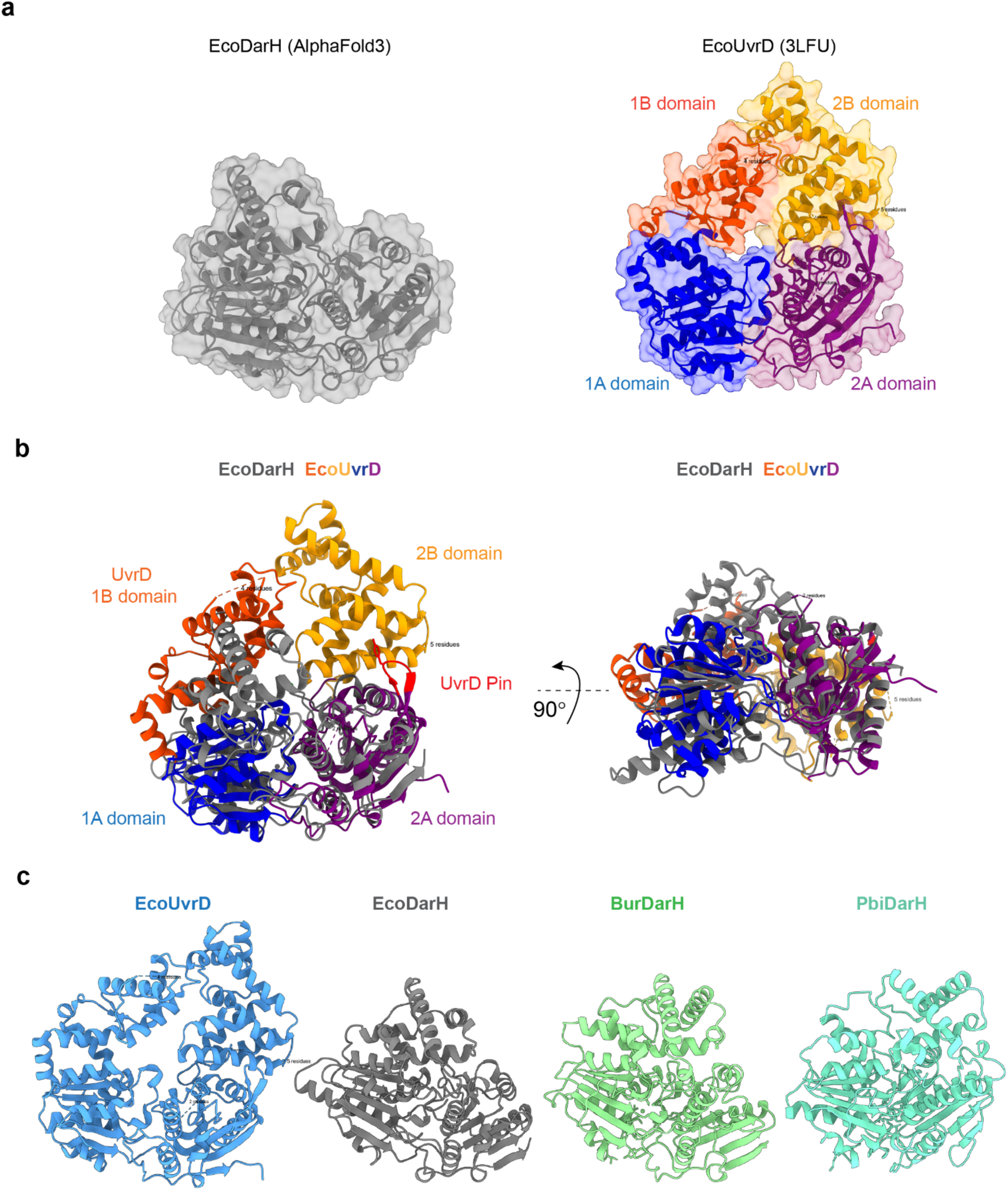
Structural comparison of DarH and *E. coli* UvrD. **a,** Predicted structure of DarH (gray), and structural model of *E. coli* UvrD, colored by domain. **b,** The same models as in **(a)**, aligned. Coloring as in (**a**), with the pin domain of UvrD colored red. Right side shows a 90 degree rotation of the model around the X axis. **c,** Structural model of UvrD (blue), or AlphaFold3 models of EcoDarH (gray), BurDarH (green), or PbiDarH (teal)

**Extended Data Figure 4:**
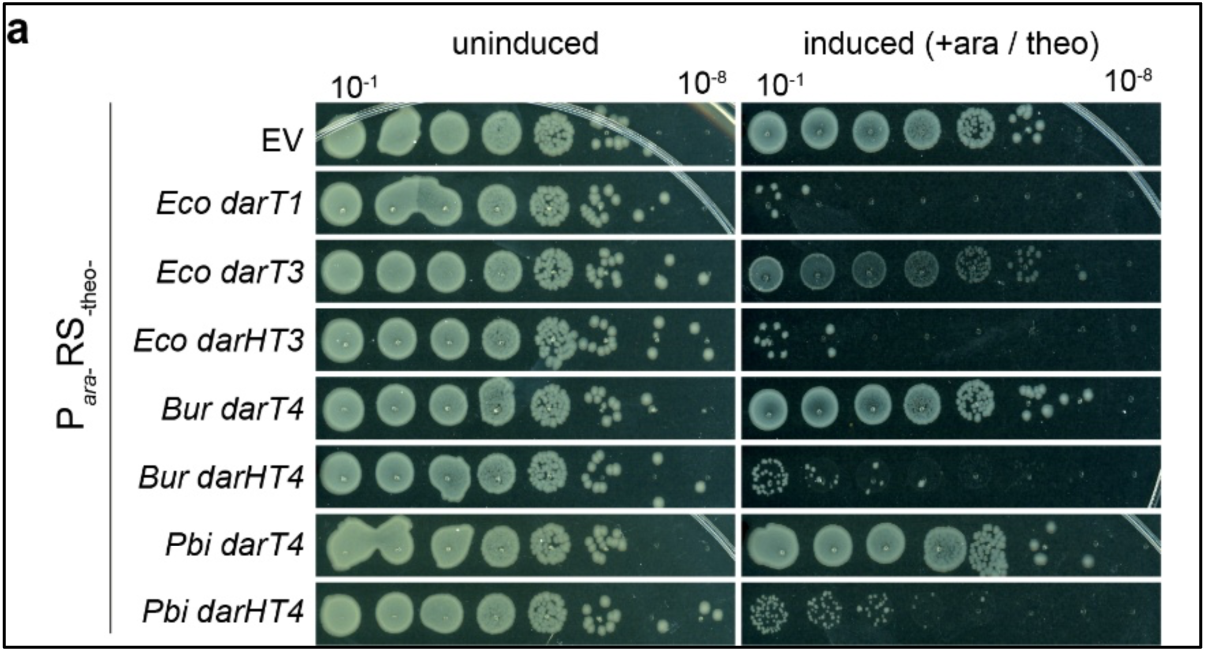
DarT3 and DarT4 require their associated DarH for toxicity. **a**, Representative CFU assay of *E. coli* BW27783 carrying the indicated *darT* or *darHT* in a P*_ara_-RS_theo_* vector. Cells were plated on LB agar alone (uninduced) or with 2mM theophylline, and 0.2%. n=3.

**Extended Data Figure 5:**
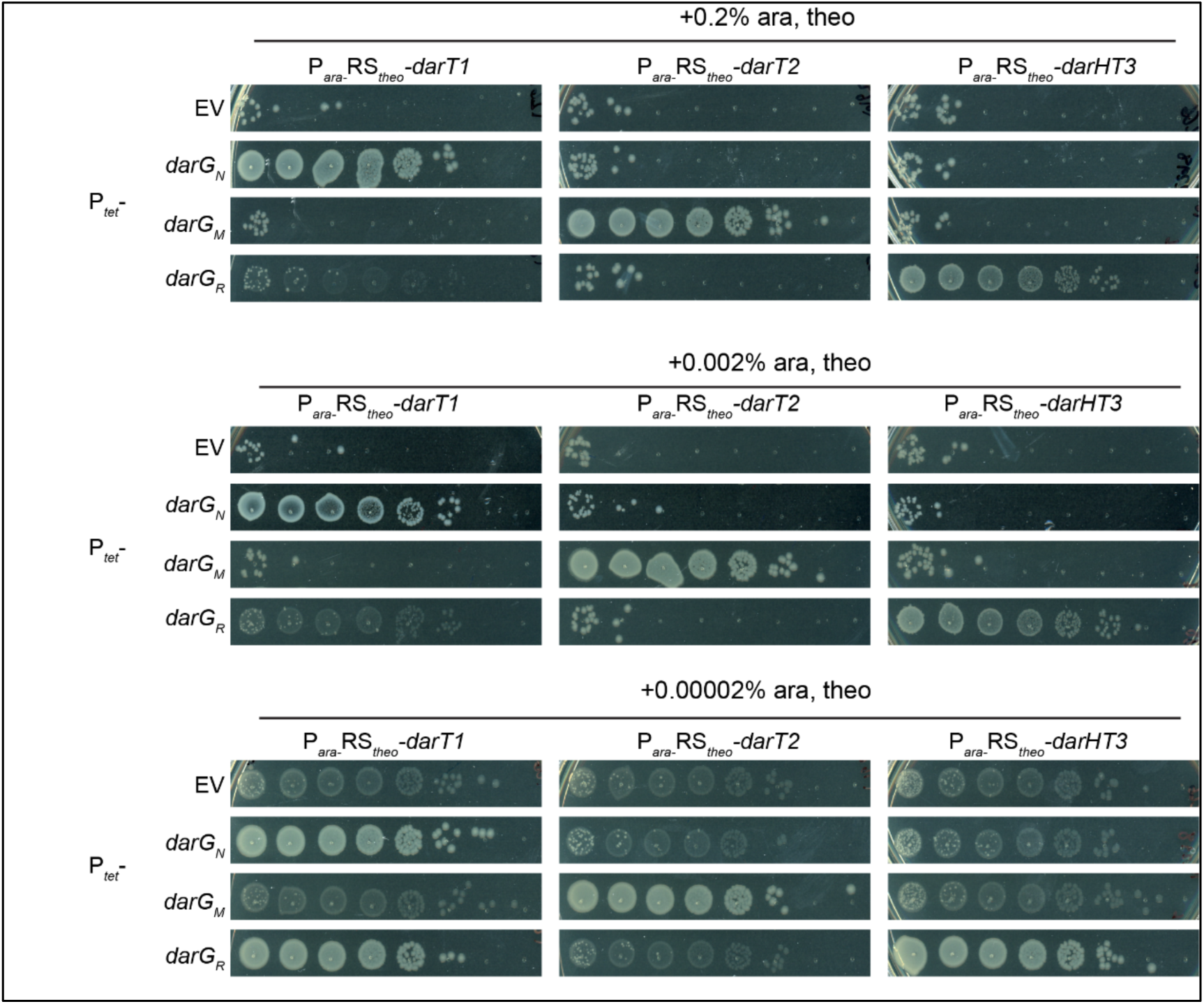
EcoDarG_R_ reverses DarT1 and DarHT3 toxicity. **a**, Representative CFU assay of *E. coli* BW27783 expressing the indicated toxin off of a P*_ara_-RS_theo_* vector, plus the indicated antitoxin off of a P*_tet_* vector. Cells were plated on LB agar with 10 ng/mL aTc, 2 mM theophylline, and 0.2% (top), 0.002% (middle), or 0.00002% (bottom) arabinose.

**Extended Data Figure 6:**
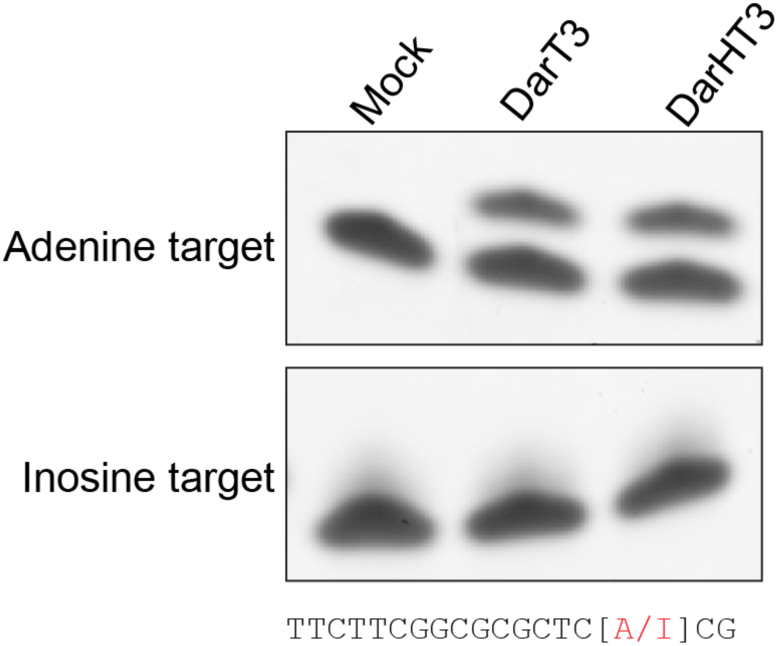
Inosine is not ADP-ribosylated by DarHT3. **a,** Urea-PAGE gel shift of an oligo with a single adenine or inosine (shown in red below), following incubation with *E. coli* DarT3 or DarHT3. n=1.

**Extended Data Figure 7:**
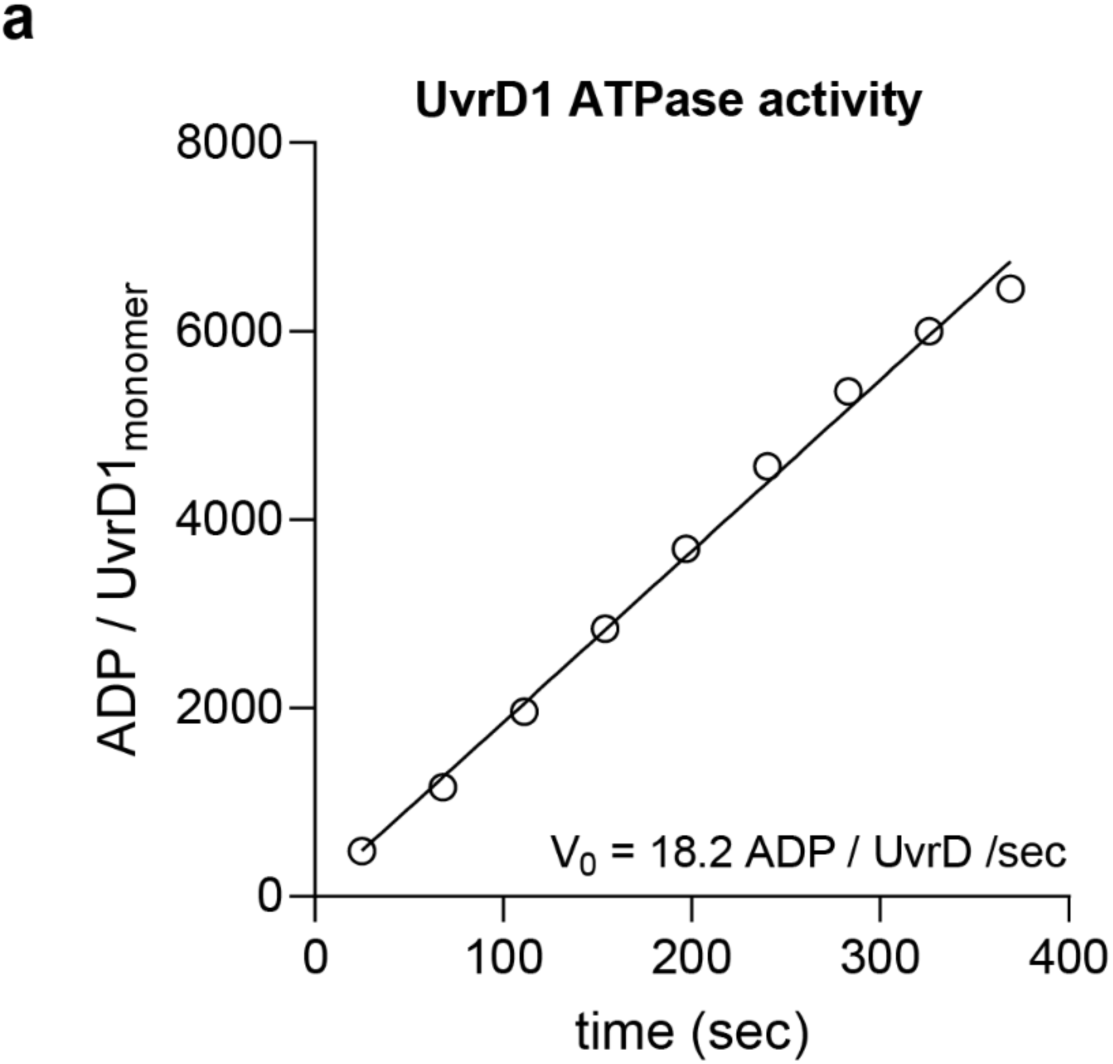
ATP hydrolysis rate of E. coli UvrD. **a,** UvrD1 (10nM) ATPase activity over time in the presence of ssDNA. Line of best fit represents a linear regression.

**Extended Data Figure 8:**
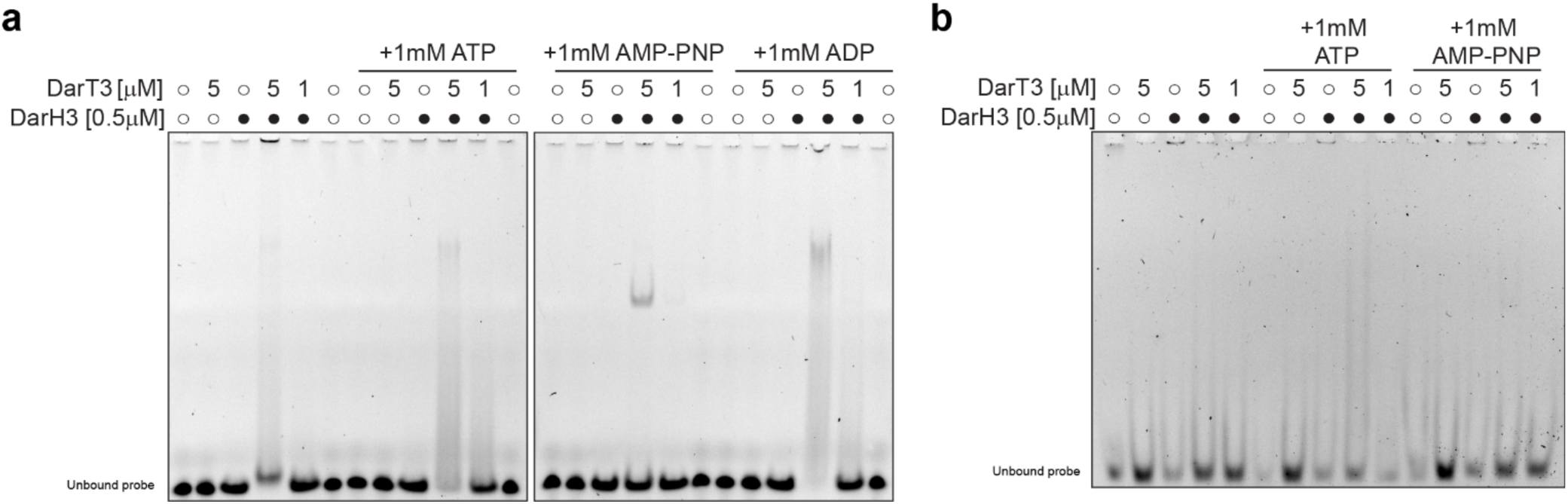
EcoDarHT3 binds to short ssDNA probes, but not dsDNA. **a,** Representative EMSA showing binding to a fluorescein-labeled 33-base ssDNA oligo. Open circles indicate absence of protein, filled circles indicate presence of DarH (0.5 μM). DarT3 concentration indicated by numbers. n=2. **b**, EMSA showing binding to a 33-base pair, blunt ended dsDNA fragment. n=2.

